# Knockout mutations of *Nicotiana benthamiana* defenses reveal the relative importance of acylsugars, nicotine, and a serine protease inhibitor in a natural setting

**DOI:** 10.1101/2024.02.26.582111

**Authors:** Boaz Negin, Fumin Wang, Hillary D. Fischer, Georg Jander

## Abstract

Plants produce an immense diversity of defensive specialized metabolites. However, despite extensive functional characterization, the relative importance of different defensive compounds is rarely examined in natural settings. Here, we compare the efficacy of three *Nicotiana benthamiana* defensive compounds, nicotine, acylsugars, and a serine protease inhibitor, by growing plants with combinations of knockout mutations in a natural setting, quantifying invertebrate interactions, and comparing relative plant performance. Among the three tested compounds, acylsugars had the greatest defensive capacity, affecting aphids, leafhoppers, spiders, and flies. Nicotine mutants displayed increased leafhopper feeding and aphid colonization. Plants lacking both nicotine and acylsugars were more susceptible to flea beetles and thrips. By contrast, knockout of the serine protease inhibitor did not affect insect herbivory in the field. Complementary experiments under controlled laboratory conditions with caterpillars grasshoppers, and aphids confirmed results obtained in a natural setting. We conclude that the three metabolite groups collectively provide broad-spectrum protection to *N. benthamiana*. However, there is a gradient in their effects on the interacting invertebrates present in the field. Furthermore, we demonstrate that, even if individual metabolites do not have a measurable defensive benefit on their own, they can have an additive effect when combined with other defensive compounds.

## Introduction

Plants synthesize an enormous diversity of specialized metabolites, with estimates ranging from 200,000 to 1,000,000 unique compounds in the plant kingdom (Wang *et al*., 2019). Any individual plant produces roughly 1,750 – 3,500 specialized metabolites with diverse functions (Pichersky & Lewinsohn, 2011). Many families of specialized metabolites, such as benzoxazinoids (Meihls *et al*., 2013), cardiac glycosides (Agrawal *et al*., 2012), and glucosinolates (Hopkins *et al*., 2009), have well-documented functions in plant defense, whereas others are yet to be investigated. All plants produce multiple classes of specialized metabolites with differing defensive functions. For instance, *Erysimum cheiranthoides* (wormseed wallflower) accumulates both cardenolides and glucosinolates (Züst *et al*., 2020) and *Nicotiana attenuata* (coyote tobacco) produces nicotine (Steppuhn *et al*., 2004), chlorogenic acid (Demkura *et al*., 2010), protease inhibitors (Zavala *et al*., 2004) and volatile terpenes (Yang *et al*., 2023), which each have a distinct protective role. Furthermore, the effects of the different defensive metabolites are not only additive, but also can involve more complex interactions. Examples include a protective function only in the presence of additional metabolites (Steppuhn & Baldwin, 2007), reciprocal reduction in toxicity (Heiling *et al*., 2022), or co-option by specialist insect herbivores for their own protection (Reichstein *et al*., 1968; Hoogshagen *et al*., 2023). Therefore, altering the production of several defensive compounds simultaneously and examining their ecological impacts in a natural setting are essential for gaining insight into their functions and interplays in plant defense.

Biosynthesis of nicotine, a pyridine alkaloid built from the conjugation of a pyridine ring and a nitrogen containing methylpyrrolidine ring (Kaminski *et al*., 2020), is restricted to *Nicotiana* species (Negin & Jander, 2023). The function of nicotine in protecting plants against insect herbivores has been extensively demonstrated under both laboratory (Yang & Guthrie, 1969; Parr & Thurston, 1972) and field conditions (Steppuhn *et al*., 2004). Nicotine functions primarily by targeting nicotinic acetylcholine receptors in animal nervous systems (Millar & Denholm, 2007), and investigations of tritrophic interactions exemplify the complex relation between this specialized metabolite, insects that consume it, and their predators (Kumar *et al*., 2014; Xu *et al*., 2017).

Acylsugars are a diverse family of compounds with the common feature of a sugar core that has several aliphatic chains of different lengths connected to it (Fan *et al*., 2019). These metabolites, which are secreted from glandular trichomes of plants in the Solanaceae and several other families (Fobes *et al*., 1985; Kroumova & Wagner, 2003; Schilmiller *et al*., 2012), have an inhibitory effect on insect herbivores in controlled insect-plant interaction studies, both *in planta* (Van Dam & Hare, 1998; Hare, 2005; Luu *et al*., 2017; Feng *et al*., 2022) and *in vitro* (Puterka *et al*., 2003). Several studies have identified acylsugar biosynthesis genes in tomato and other Solanaceae (Schilmiller *et al*., 2012, 2016; Lou *et al*., 2021), paving the way for targeted manipulation of these defenses.

Protease inhibitors are proteins that inhibit insect digestive enzymes, rendering the consumed plant material less nutritious (Green & Ryan, 1972). These plant proteins, which are induced in response to caterpillar feeding, jasmonate treatment, or wounding (Van Dam *et al*., 2001), reduce caterpillar growth (Zavala *et al*., 2004; Hu *et al*., 2018), slow aphid development (Carrillo *et al*., 2011), and reduce attractiveness of plants to mirids (Zavala *et al*., 2004). Field-grown *Solanum nigrum* (black nightshade), in which expression of three serine protease inhibitors was silenced, was more susceptible to both thrips and caterpillars (Hartl *et al*., 2010).

*Nicotiana benthamiana* is an Australian tobacco species that adapted to desert conditions millions of years ago (Bally *et al*., 2018). This species is widely used for plant molecular biology research, largely due to its extreme viral susceptibility (Jassbi *et al*., 2017), amenability for transient gene expression assays (Bally *et al*., 2015), and ease of heritable gene editing (Ellison *et al*., 2020). By contrast, very few laboratory studies involve the investigation of insect interactions in this species, likely due to the high level of endogenous herbivore resistance.

In the current study, we utilized existing genome-edited *N. benthamiana* plants, newly-generated mutants, and mutant combinations to examine the function of three representative defensive compounds, nicotine, acylsugars, and a serine protease inhibitor, in both a natural setting and under controlled laboratory conditions. We found, that acylsugars had the strongest effect, nicotine had a comparatively moderate effect, and the protease inhibitor had no discernible effect on invertebrate interactions.

## Materials and Methods

### Plant material and growth conditions

*Nicotiana benthamiana* plants were grown in temperature-controlled growth chambers (Conviron, Winnipeg, Canada) at 28°C or in a temperature-regulated greenhouse, at ∼27°C for approximately three weeks prior to transplanting in the field. For laboratory insect assays, *N. benthamiana* plants were grown similarly in growth chambers at 28°C – 30°C. Plants used for genome editing either expressed the *PCO-CAS9* gene (Li *et al*., 2013), or the same *CAS9* while also harboring a mutation in the *SerPIN-II* gene (Feng & Jander, 2023) described as the *SERP-II3-1* line (see Table S1 where this line, harboring a three BP deletion is shown to be named *serp2-2* in the current paper). These were grown in a growth room at 22°C prior to and following viral infection. For assessment of the effects of growth temperature, on plant size, plants were grown in a 22°C growth room, or in growth chambers at 28°C or 34°C.

Field-grown plants were planted during the summer of 2023 in a deer-excluded field in Ithaca, NY (42.46 °N, 76.44 °W). Plants were planted in blocks of 14, each including two wildtype plants and two independent lines of each mutant combination (Fig. S1). The plants’ locations in the blocks were randomized, but changes were made to ensure each mutation was equally represented in the inner plants of the block and in the different compass directions. Distances between individual plants were 40 cm and 110 cm between blocks (building on the field experimental setup of Steppuhn et al (Steppuhn *et al*., 2004)), and 24 blocks were planted in three rows (Fig. S1B).

### Insects and growth conditions

*Chloridea virescens* (tobacco budworm; formerly *Heliothis virescens*) and *Spodoptera exigua* (beet armyworm) eggs were purchased from Benzon Research (Carlisle, PA, USA) and hatched on beet armyworm diet (Southland Products, Lake Village, AR, USA) in a 28°C incubator. Two strains of green peach aphid (*Myzus persicae*) were used in this study: a green strain that was reported to be nicotine-sensitive and a red “USDA” strain that is nicotine adapted (Ramsey *et al*., 2007, 2014). The green strain was reared on *Brassica rapa* var chinensis, whereas the red strain was reared on *Nicotiana tabacum*. *Schistocerca americana* (American bird grasshopper) eggs were kindly provided by Hojun Song at Texas A&M University. Eggs were hatched in a 30°C growth chamber, and nymphs were raised at 30°C on lettuce, wheat grass, and wheat bran.

### Mutant accessions and generation of the mutant population

The mutant population used in this study consisted of wildtype plants, mutants of three genes encoding defensive compounds, and their combinations: *SerPIN-II* (Feng & Jander, 2023) (the *serp2-1* line corresponding to the *serpII3-2* line and the *serp2-2* line corresponding to *serpII3-1*), *ACYLSUGAR ACYL TRANSFERASE2* (*ASAT2*;(Feng *et al*., 2022)), and the *N. benthamiana A622* gene, which was identified by a BLAST search of the *N. benthamiana* transcriptome with known genes from other *Nicotiana* species (Deboer *et al*., 2009) and was predicted to convert nicotinic acid into 3,4-dihydronicotinic acid as an intermediate in nicotine biosynthesis. The *N. benthamiana* gene with the highest similarity to the *A622* query sequence was Nbe.v1.s00140g13970 (Supplemental Data S3). Three crRNA (RNA sequences transcribed from the CRISPR locus) targets (TCTCTGATAAGAACGAAAGT, TGATTTCCACTGTCGGGGGA, and AGAGTCTTATTCAATGTCCG) in the first, second and fourth exons, respectively, of *A622* were identified using the CRISPR-P tool (Lei *et al*., 2014). The three crRNAs were inserted into a tobacco rattle virus (TRV) plasmid vector (Ellison *et al*., 2020), either on their own or in combination with an additional two crRNAs targeting *ASAT2*, which were identical to those used by Feng et al., (Feng *et al*., 2022). The two guide RNA combinations were expressed on two different backgrounds, either *CAS9*-expressing plants (Ellison *et al*., 2020) or *CAS9*-expressing *SerPIN-II* mutants (Feng & Jander, 2023).

Genome editing of *N. benthamiana* plants was performed using TRV expressing guide RNAs (Ellison *et al*., 2020). In short, the PEE392 plasmid (Addgene #149282, www.addgene.org) was used as a template to amplify a guide RNA fragment, a changing crRNA and a fragment from the *FLOWERING LOCUS T* (*FT)* gene to drive meristem localization. PCR products were run on electrophoreses gels, cut, cleaned and combined into the PEE083 plasmid (Addgene #149279) using the Golden Gate assembly system (Engler *et al*., 2008). PEE083 is a TRV plasmid encoding TRV2 coat proteins, and between them the guide RNAs driven by a yellow leaf curl virus promoter. This plasmid was transferred into the GV3101 *Agrobacterium* strain and co-infiltrated with a TRV1 vector to drive viral infection of *N. benthamiana* (Sparkes *et al*., 2006). Plants were then grown to seed set, younger seed capsules were collected, and these were grown for analysis by liquid chromatography-mass spectrometry (LC-MS).

### Identification and relative quantification of nicotine and acylsugars

Nicotine and acylsugars were identified and relatively quantified using LC-MS. Acylsugars were extracted using a 3:2:2 mix of acetonitrile (ACN), isopropanol, and water with 0.1% formic acid. Leaves from the shoot apex were dipped in 500 µl of the solution and gently shaken for 2 min, following which the leaves were removed, dried, and weighed. The solvent was then filtered and analyzed using an Ultimate 3000 HPLC (Thermofisher Scientific) coupled to a Q Exactive quadrupole - orbitrap mass spectrometer (Thermofisher Scientific) using a Supelco C18 column (Sigma Aldrich; St. Louis, MO, USA). Buffers A and B were water and ACN respectively, to each of which 0.1% formic acid was added. Buffer gradients and program settings were those used in Feng et al. (Feng *et al*., 2022). For nicotine extraction, 10 mm diameter leaf discs were cut from leaves 3-4 of ∼4-week-old plants and flash frozen in liquid N_2_. Samples were ground using a GenoGrinder 1600 MiniG homogenizer (SPEX SamplePrep, Metuchen NJ, USA) and metabolites were extracted in a 500 µL solution composed of 89.9% MeOH, 10% water, and 0.1% FA. This extract was filtered and run on the same LC-MS system and column as the acylsugars. The column oven was set to 35°C, and initial flow was set to 0.1 ml/min at 99% water and 1% ACN. From 5 - 8 min, the ACN concentration was increased to 100%. At that point, the pump’s speed was increased to 0.5 ml/min and the ACN concentration was kept at 100% until 9 min, then reduced back to the initial 1% at 10 min, and held for an additional minute. Nicotine was identified using a nicotine standard (Sigma Aldrich), exact mass, and fragmentation patterns.

### Field observations

Interactions with insects, snails, slugs, and spiders were observed starting four days post planting in the field, during the morning and early afternoon hours. Due to the large number of plants initially observed, the observation cycle was divided to several days during which all the plants in the field were examined. This examination included observation of plants without disturbing them, from a distance of ∼20 cm. Following this, interactions were examined more closely on both sides of the leaves, the shoot apexes and the stems. For identification, interactors were photographed using a Cannon G16 camera, or were sampled and further examined under a stereo microscope. All 336 plants were examined between days 4-7 and 18-23. Following this, the first 8 blocks, which were severely damaged in an uneven manner, were no longer examined for invertebrate interactions. The remaining 16 blocks were examined between days 31-34, and blocks 10-13 and 18-22 were examined on day 45. Finally, blocks 12, 18 and 20 were examined on day 59. Given the two-week period between observations and the extreme mobility of most interactors, invertebrates that were observed during following visitations were scored individually. This approach was validated by an analysis of observations of prominent invertebrates (leaf hoppers, aphid colonies, flea beetles and spiders; Supplemental Data S2) in relation to the specific plants on which they were found, which showed that in the vast majority of cases these insects were observed on plants that did not have similar insects on them in the previous observation, did not have similar insects in the following observation, and there was no case of insect types observed continually on any plant. Furthermore, even when invertebrates were found on the same plant on consecutive observations, they were at times from different species altogether (Fig. S2). In addition to invertebrate interactions, plant height and survival were monitored. Plant height was recorded at 24, 36-38, 52 and 66 days. Survival was monitored continually, *i.e.*, when a dead plant was identified, it was recorded as such and added to the analysis.

Field grown plants were also sampled for nicotine and acylsugars. This process was similar to sampling of these metabolites in the lab except that nicotine samples were whole young leaves from the shoot apex and were weighed.

### Insect assays under controlled conditions

For caterpillar growth assays, freshly hatched *S. exigua* neonates were placed in organza mesh bags (Amazon item B073J4RS9C) on individual *N. benthamiana* leaves and returned to a 28°C growth chamber. After 11 days, survival was scored, and live caterpillars were weighed. This timeframe was chosen following a previous trial with *Nicotiana glauca* in which *S. exigua* larvae were confined on leaves for a similar duration (Negin *et al*., 2024). Four-day-old *C. virescens* larvae hatched on beet armyworm diet were weighed, and those weighing 5-12 mg were placed in organza mesh bags on individual *N. benthamiana* leaves, with a single caterpillar in each bag. Plants with caterpillars were then moved back to 28°C growth chambers. After four days, the caterpillars were weighed, and survival was assessed. Relative growth rate (RGR) was calculated using the equation 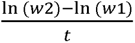 where *w1* is the weight on day 4, *w2* is the weight on day 8, and *t* is the number of days between weighing (4 days in this case). This equation represents how many mg the caterpillars gained every day for every mg they weighed.

Seven-week-old wildtype, *asat2*, a*622*, and *asat2+a622 N. benthamiana* plants, grown at 30°C so that all plants are similarly-sized, were used for *Schistocerca americana* choice and no-choice assays. For no-choice assays, four two-week-old grasshoppers were confined on individual *N. benthamiana* plants using 10 x 16 inch micro-perforated polypropylene bags (PrismPak; Berwick, PA, USA). Bagged plants with grasshoppers were placed in an incubator at 30°C. After 16 days, grasshopper survival was assessed and the surviving grasshoppers were weighed. For choice assays a cork borer was used to make 23 mm diameter leaf disks from mature *N. benthamiana* leaves. Individual leaf disks of the two genotypes being compared were placed on moistened filter paper in 10 cm diameter Petri dishes. Two two-week-old grasshoppers were added to each Petri dish, and the Petri dishes were placed in a growth room at 23°C. After 14 hours, the leaf material was collected and scanned. The area of the remaining leaf material was calculated using ImageJ (https://imagej.net/ij/). To account for leaf shrinkage during the experiment, the amount eaten by the grasshoppers was calculated relative to leaves that had received no grasshopper feeding.

Aphid reproduction assays were performed using both nicotine-sensitive and adapted *M. persicae* strains. Five adult aphids were caged on the adaxial side of an intact leaf for parthenogenetic reproduction. After 24 h, adults were removed and only three newly born nymphs were left in each cage. After either eight or nine days (for nicotine-sensitive and adapted strains, respectively), aphids were counted. Leaf cages were placed on individual plants in blocks of all seven genotypes used in this study (wildtype, *a622*, *asat2*, *serp2*, *a622*+*asat2*, *a622*+*serp* and *a622*+*asat2*+*serp2*). For choice assays, two leaf discs from different genotypes were placed on top of Whatman filter paper in 100×15 mm Petri dishes, to which 1.1 ml water were added. Five grown aphids were placed between the two discs and the Petri dishes were sealed with Parafilm. These were left in a growth room for 24 h, after which the number of aphids on each of the leaf discs was counted.

### Statistical analysis

Statistical analyses were performed using JMP 16 software (SAS Institute, Cary, NC, USA). The frequency of field observed events was analyzed using a Chi-square test comparing the different proportion of plants with a specific invertebrate observed on them between mutants and wildtype. A Benjamini-Hochberg correction was performed to account for multiple comparisons. In this analysis, events were considered as a type of invertebrate on a plant (*i.e*., a plant with 3 leafhoppers was scored as one leafhopper event). Survival was similarly assessed using a Chi-square test, whereas plant heights and field extracted nicotine and acylsugars were compared using a Tukey’s HSD test. Caterpillar and grasshopper weights on the different mutants were first compared using a two-way ANOVA to examine the effects of the different metabolites and the interactions between them. Next, caterpillar weights on individual mutants were compared to those on wildtype leaves using Student’s *t*-tests followed by a Benjamini-Hochberg correction. This approach was chosen to balance a conservative approach, which would account for multiple comparisons, but would not lose power through unnecessary comparisons (such as comparing *serp2* to *a622*+*asat2*), given the inherently large variance in insect assays. Aphid reproduction was analyzed similarly. For choice assays, paired Student’s *t*-tests were used.

## Results

### Generation of a622 and asat2 mutants

To generate *N. benthamiana* lines with reduced nicotine and acyl sugar content, the *A622* gene, which is key to nicotine production (Deboer *et al*., 2009), and *ASAT2*, which is essential for acyl sugar production (Fan *et al*., 2016), were targeted for CRISPR mutagenesis. Plants were infected with tobacco rattle virus (TRV) carrying guide RNAs (gRNAs) targeting either the *A622* gene, or a combination of the *A622* and *ASAT2* genes in both *CAS9*-expressing wildtype and *CAS9*-expressing *serp2* mutants, which do not produce the Serp-II3 protease inhibitor (Feng & Jander, 2023). We assessed foliar nicotine and acylsugar content in subsequent generations using LC-MS (Fig. S3-S4). Multiple independent lines from both backgrounds and infiltrated constructs contained less than 5% of normal nicotine abundance (Fig. S3). When analyzing acylsugar abundance, we found plants containing less than 5% of wildtype levels of acylsugars in the *CAS9* background (Fig. S3D), but no plants with less than 5% of the acylsugar abundance in the *serp2* background (Fig. S3F). However, there was one line close to the 5% threshold whose progeny were grown for further examination.

When confirming the mutations using Sanger sequencing, we found that the *A622* gene was often missing stretches of DNA between the three gRNA targets (1-3, 1-2, or 2-3), but single-base pair indels were also formed (Fig. S5, Table S1). The *ASAT2* gene displayed a range of mutations, including 136, 29, 3, and 2-base pair deletions (Fig. S6, Table S1). Based on these results, additional plants were grown and screened (Fig. S3, S4) until the final panel for field experiments was assembled. This panel did not include the *serp2*+*a622*+*asat2* triple mutant, which was still being characterized and would be included later in controlled laboratory experiments. Furthermore, when validating genome edits in the acylsugars + serpin double mutants, both independent lines were not edited in the *ASAT2* gene. These plants were excluded from further analyses.

### Effects of nicotine, acylsugars and SERP2 on invertebrate interactions in a natural setting

Prior to transferring the mutant population to the field, we noticed that plants harboring mutations in the *A622* gene were smaller than wildtype and the other *serp2* and *asat2* mutants. This phenotype has been previously reported in field grown *Nicotiana tabacum*. However, we found that this growth retardation was variable, with plants from the same genotype sometimes exhibiting growth retardation and sometimes not. When searching for conditions that would facilitate normal growth of the *a622* mutants, we found that growing the plants in higher temperatures in growth chambers or a greenhouse reduced the growth retardation significantly (Fig. S7). Whereas *a622* mutants grown at 22°C were significantly smaller than wildtype, those grown at both 28°C and 34°C were similar in size to wildtype. Examining the response to the changes in the growth temperature using a two-way ANOVA showed a significant interaction between the line and growth temperature, meaning that wildtype and *a622* responded differently to the increase in temperature. Wildtype plants were smaller as the temperature increased. By contrast, *a622* plants were larger at 28°C than at 22°C, but slightly smaller at 34°C than when grown at 28°C. Therefore, plants were grown in a temperature-controlled greenhouse at 27°C or in growth chambers at 28°C. Wildtype *a622,* and all genotypes of the mutant population, which showed similar growth under these conditions, were planted in the field.

The mutant population was planted in 24 blocks of 14 plants in a plowed 20 x 10 m field section (Fig. S1). Invertebrate interactions were observed at five time points up to day 59 post-planting (Fig. 1, S8; Supplemental Data S1). Throughout the wet summer months, snails and slugs caused the greatest amount of plant damage. During the first observation cycle (4-7 d post-planting) many leafhoppers and flies but no aphid colonies were observed. During the second cycle (18-23 d post planting), six aphid colonies, fewer flies, and more than double the previous number of flea beetles were observed. During the third observation cycle (31-34 d post-planting), aphids and aphid colonies tripled, despite fewer plants being monitored due to high mortality in the first row closest to the field margins. Far fewer leafhoppers and flea beetles were observed during this cycle, but spiders tripled. A new predator, an assassin bug, appeared on eight plants, totaling 10 individuals. At this point bumble bees pollinated plants, which had substantial numbers of flowers. During the fourth observation cycle (45 d post planting), fewer plants were monitored. However, aphids and aphid colony numbers were maintained whereas other herbivore numbers were reduced. Assassin bug numbers increased to 29 individuals, whereas spiders maintained their numbers. An additional observed interaction was four instances of aphids parasitized by wasps. Finally, the fifth observation cycle (59 d post planting) included only three blocks. Despite this small number, six parasitized aphids were observed, as well as five plants with whiteflies, which were not seen in the first four observation cycles.

**Figure 1.**
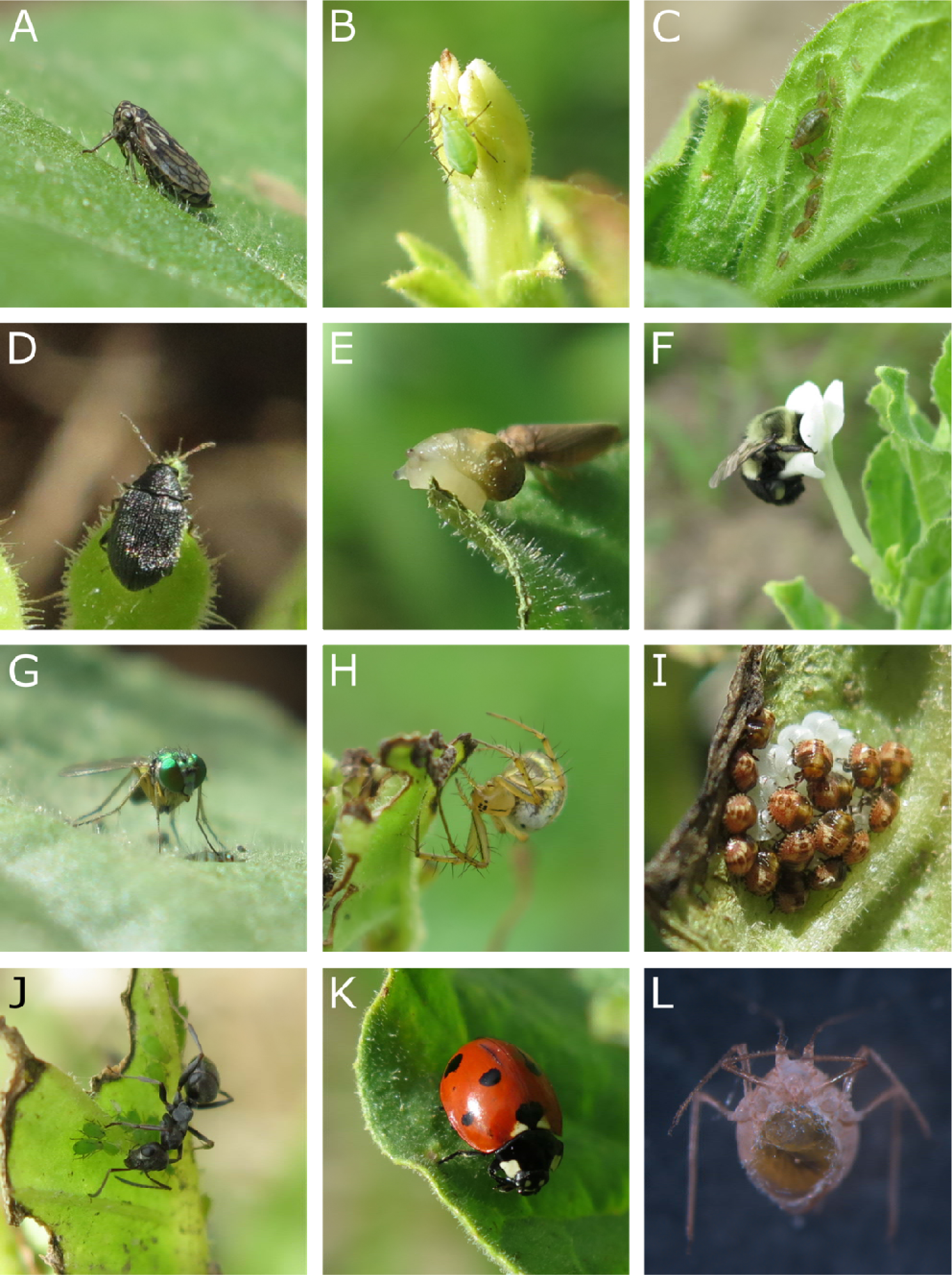
Photographs of invertebrates comprising the main interactor classes observed in the field. (A) A leafhopper from the genus *Agallia*. (B) A leaf aphid (C) A leaf aphid colony (D) A flea beetle from the *Epitrix* genus. (E). A snail, probably from the genus *Succinea*. (F) A bumble bee (genus *Bombus*) pollinating a *N. benthamiana* flower. (G) A fly from the genus *Condylostylus*. (H) An orb weaver spider, probably from the species *Mangora acalypha*. (I) Stink bug (*Halyomorpha halys*) eggs during hatching. (J) An ant “farming” an aphid colony. (K) A lady bug (*Coccinella septempunctata*). (L) A parasitized aphid collected from *N. benthamiana* with a wasp larva that has almost entirely consumed the aphid’s body.

When analyzing all of the collected data, which included over 430 individual events totaling over 560 individual invertebrates observed on plants (Fig. 2A), we discerned clear effects of some of the mutations. Leafhoppers, by far the most commonly observed insects (121 events totaling 174 individual insects), were found more frequently on *asat2*, *a622*, and *a622+asat2* mutants (Fig. 2B), demonstrating that both nicotine and acylsugars protect plants from leafhoppers (Fig. S9A). Aphids (52 events totaling 80 individuals) were found on significantly more *asat2* and *a622+asat2* mutants (Fig. 2B), indicating that acylsugars but not nicotine are important for protection against these phloem feeders (Fig. S9B). By contrast, aphid colonies were found more often on *asat2*, *a622*, and *a622+asat2* plants, and none were found on wildtype plants (Fig. 2B), indicating that despite not affecting individual aphids, nicotine does affect aphids’ ability to colonize *N. benthamiana* (Fig S9C). Flea beetles, the most abundantly observed leaf chewers (45 events, 52 individuals; Fig. 2C), were significantly more abundant on *a622+asat2* mutants, indicating that both acylsugars and nicotine provide protection, and plants only become susceptible when losing both defenses (Fig. S9D). Similar results were obtained for thrips, which were found more commonly on *a622+asat2* mutants. Slugs and snails showed a trend of being found on more *asat2* mutants. Bumble bee pollinator visitations were statistically similar between all plants due to the low numbers observed (Fig. 2C), However, the bees seemed to prefer plants that had more flowers regardless of their chemotype. In non-feeding interactions (Fig. 2D), flies and spiders preferred *asat2* and *a622+asat2* mutants.

**Figure 2.**
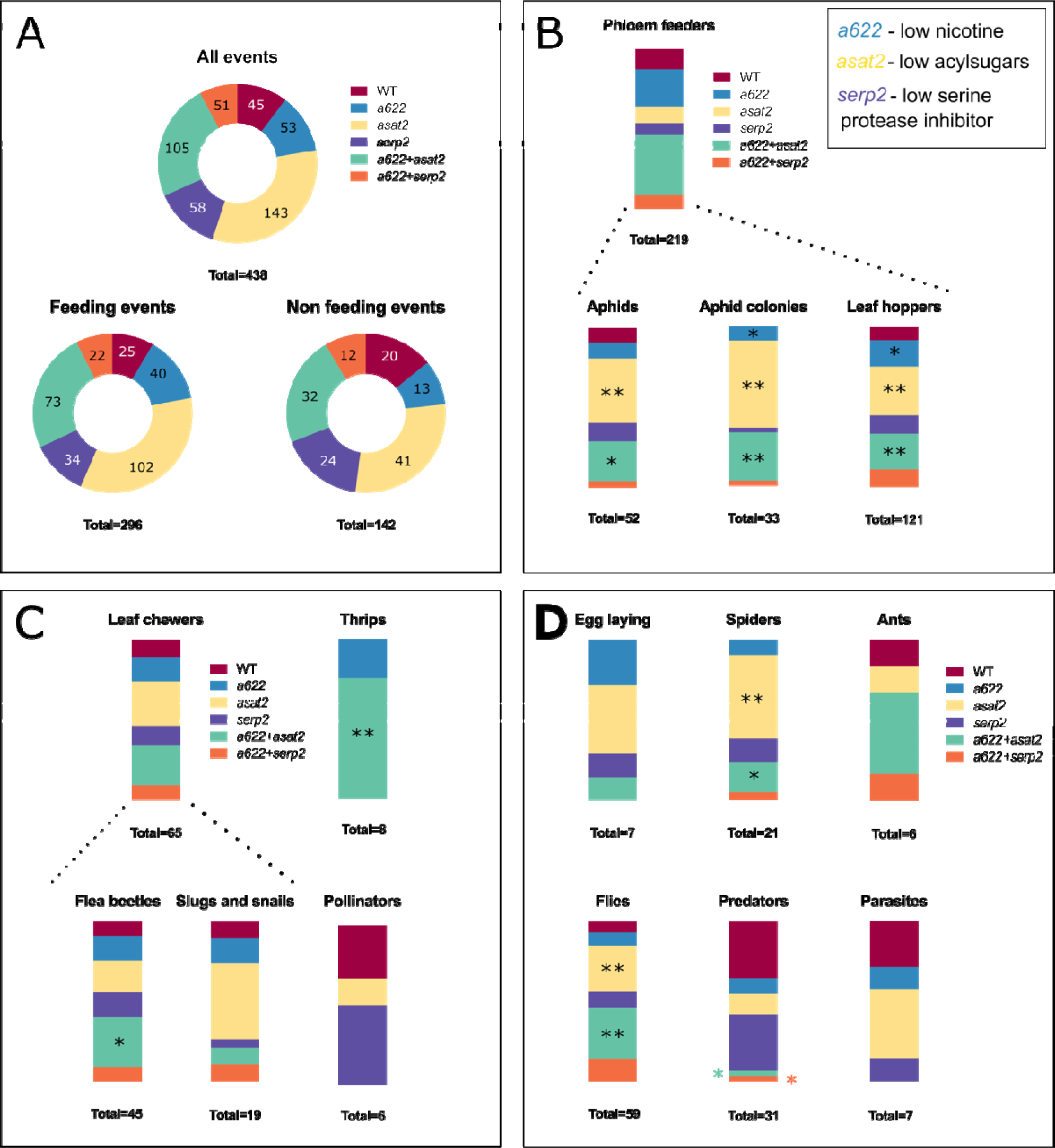
A breakdown of invertebrate interactions observed in a natural setting. (A) All the observed events separated to feeding and non-feeding interactions. Feeding interactions are further divided to (B) phloem feeders and (C) leaf chewers, with thrips and pollinators not falling strictly in either of those categories. (D) Non-feeding interactions. These include all interactions in which invertebrates interacted with the plant but did not directly feed on it. Within each invertebrate type, frequencies were compared between plant treatments and wildtype by a Chi-square test, as proportions of plants with events out of all the observed plants of the genotype and followed by a Benjamini-Hochberg correction (*p<0.05, **p<0.01). Interactions which were observed at very low frequencies, such as a single caterpillar, were not included as individual bars, but are accounted for in their group’s proportion (such as leaf chewers). Wildtype (WT) n = 132, *a622* n = 107, *asat2* n = 131, *serp2* n = 132, *a622* + *asat2* n = 107, *a622* + *serp2* n = 101.

Insect predators, including lady bugs and assassin bugs, were found significantly less commonly on *a622+asat* and *a622+serp* plants. However, this is most likely due to the different plant sizes, as plant height at the end of the experiment correlates significantly with average assassin bug abundance on the different chemotypes (R^2^=0.83, p=0.0042). Three other observation classes, ants, egg-laying, and parasites, did not show differences between the mutants due to their rare occurrence (6-10 events). However, it is interesting to note that the ratio of parasitized aphids to aphids and aphid colony occurrence, is not correlated (R^2^=0.17, p=0.35 and R^2^=0.04, p=0.68 respectively). This lack of correlation could indicate that wasps parasitize aphids with greater ease when aphids are on better-protected plants.

We next examined whether *asat2*, *a622*, and *a622+asat2* mutants were significantly different from each other, indicating that effects seen in double mutants were additive, and whether one of these specialized metabolites had a stronger effect than the other (Fig. S9). We compared seven interactions in which some of these mutants were significantly different from wildtype. There was no difference between the three mutants in leafhopper, flea beetle, and thrips interactions, indicating that both nicotine and acylsugars protect plants similarly from these insects. In the case of aphids, aphid colonies, and flies, both *asat2* and the double mutants were significantly different from nicotine mutants, indicating that only acylsugars protect plants from aphids and flies and that their protection from aphid colonization is stronger than nicotine, to the extent that nicotine has no added effect on this interaction when acylsugars are affected. Spiders were affected by acylsugars and the double mutants, indicating an effect of acylsugars but not nicotine on these invertebrates.

When examining general fitness parameters, we monitored plant height and survival. In general, all lines that harbored mutations in the *A622* gene were smaller (Fig. 3C). However, this is an *a622*-specific phenotype, which also can be seen when growing *a622* mutants under controlled conditions (Fig. S7). Since we found that cold temperatures contributed to the dwarfed phenotype, it may very well be that the cold summer in the field in Ithaca, NY, contributed to *a622* mutants growing more slowly, and it is difficult to attribute this phenotype to the effect of invertebrate interactors. In addition to the effects of the *a622* gene, at the last measurement point *asat2* mutants were significantly smaller than wildtype, indicating a cumulative effect of their increased invertebrate susceptibility. *Serp2* plants were similar to wildtype at all time points. In addition to height, plant survival was assessed (Fig. 3D). Here we found that all plants harboring the *a622* mutation survived at lower rates, whereas *asat2* mutations did not affect plant survival. Seed production could not be assessed because, due to the transgenic status of the plants, capsule development was terminated at early stages. However, assuming that the plants’ harvest index was not drastically altered, we can assume that the smaller biomass of *asat2* mutants would correlate to a reduced seed yield.

**Figure 3.**
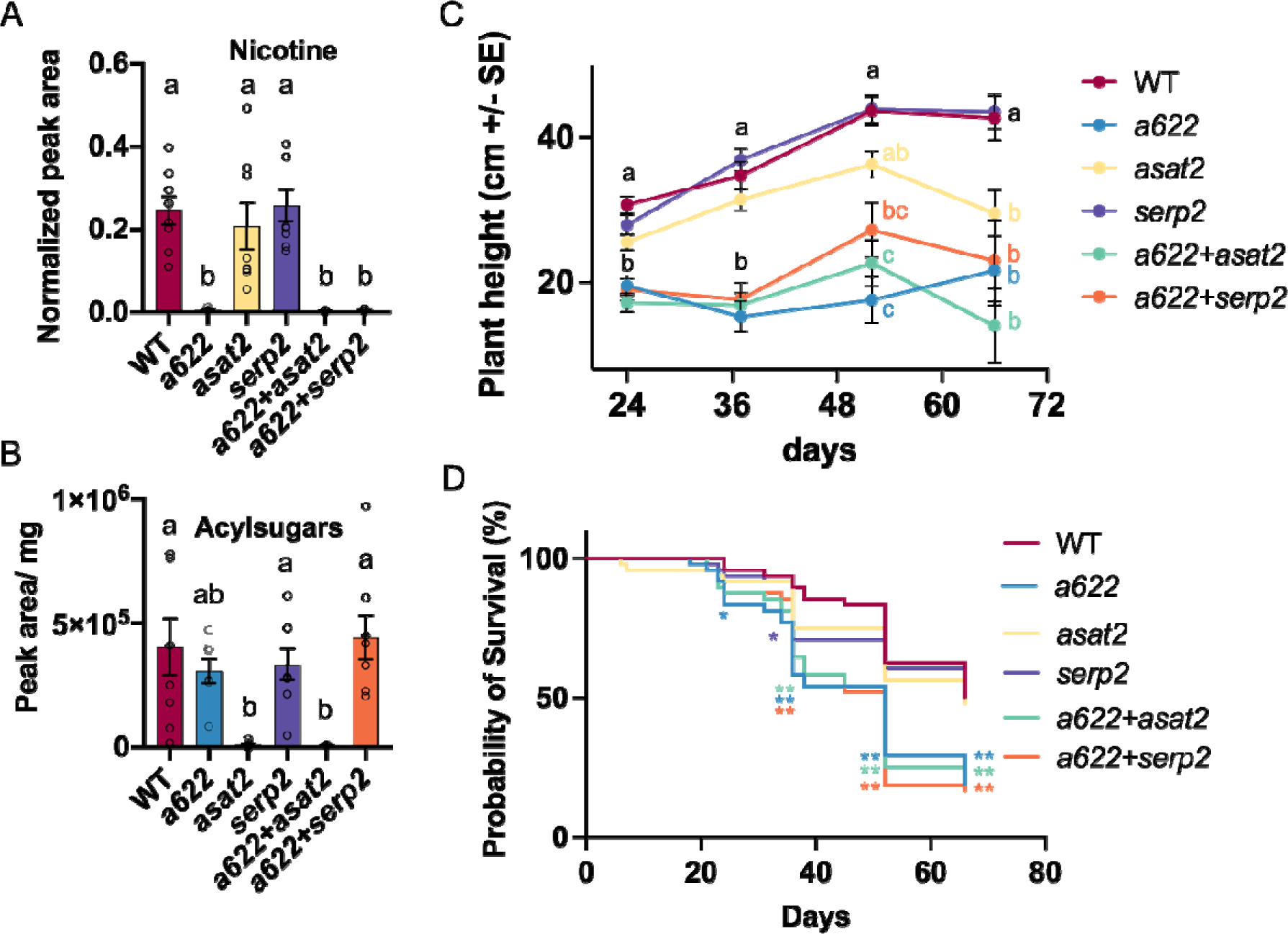
Nicotine and acylsugar abundance of field-grown plants, plant height, and survival. (A) Nicotine abundance determined by LC-MS with peak normalization by a bipyridine internal standard and tissue weight. (B) Acylsugar abundance determined by LC-MS with the peak area normalized to tissue weight. (C) Height of field grown plants throughout the growing season. Dead plants (as scored by lack of stem greenness) were removed from the analysis. Different letters indicate significance of p < 0.05 as determined by Tukey’s HSD test. Results in A-C are presented as the mean ± SE. (D) Survival frequency of field grown *N. benthamiana*. *p < 0.05 **p < 0.01. as determined in a Chi-square test, comparing wildtype and each mutant. (A-B) N = 7-8, (C) 24 d n = 40-46, 37 d n = 26-41, 52 d n = 11-30, 66 d n = 8-26. (D) N = 48.

### Insect interactions under controlled conditions

Interactions observed in the field are the sum of many different factors combined into a single phenotype. Therefore, we next examined the two main feeding guilds, phloem feeding and leaf chewing, under controlled conditions. These experiments, in which plants were size-matched, also aided us in understanding whether the smaller frequency of interactors found on *a622* mutants compared to *asat2* mutants in the field was solely due to their smaller size. We initially examined the interaction of the broad generalist caterpillar *Spodoptera exigua* (beet armyworm) using a population that also included the *a622+asat2+serp2* triple mutant. However, only 10 caterpillars out of 350 neonates placed on 179 plants from the seven genotypes survived, and no trend was seen of higher survival on plants that had fewer defensive compounds. Therefore, we performed an additional trial with *Chloridea virescens* (tobacco budworm), with caterpillars 4 days post-hatching, rather than first-instar larvae. Caterpillar weight (5 – 12 mg) was recorded and insects were placed in mesh bags on individual leaves. Following four days of feeding, the caterpillars reached weights of 4.5-108 mg. the average weight per treatment varied between 46.4 mg for wildtype and 59.8 mg for the triple mutants. Of the seven mutant combinations examined, only the triple *a622*+*asat2*+*serp2* mutants had a significantly higher relative growth rate (RGR) than wildtype (Fig. 4A). This result is intriguing since the *serp2* mutants showed no effect in field and controlled conditions, but the addition of the *serp2* mutation beyond the *a622*+*asat2* mutations lead to caterpillars being heavier than on wildtype. When the effects of the three defensive compounds on the caterpillars’ RGR were examined in a two-way ANOVA, a significant effect was found only for the *asat2* mutation (p = 0.0079), with no significant effect for the interactions of the compounds, indicating that acylsugars affected *C. virescence* similarly when only they were lacking and when combined with the other compounds. In this instance, only 13 out of 337 caterpillars placed on the plants were dead or had escaped at the end of the experiment, with no trend seen for more death and escape on any of the examined genotypes.

**Figure 4.**
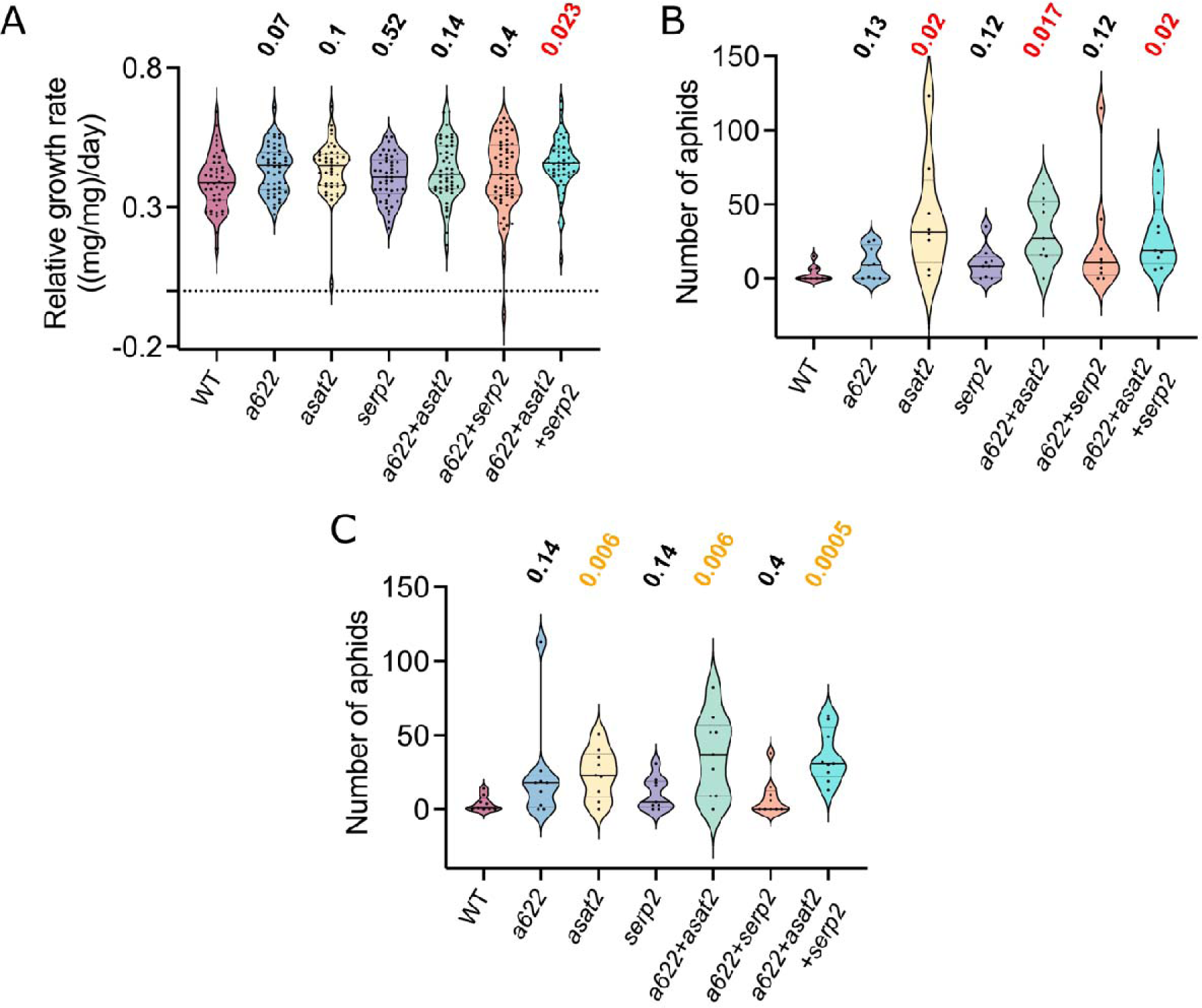
*Chloridea virescens* growth and *Myzus persicae* reproduction under controlled conditions. (A) *C. virescens* weight following four days of feeding on *N. benthamiana* leaves. Larvae were placed in mesh bags on individual leaves four days post-hatching. N = 44-54. (B-C) The number of aphids in leaf cages, 8 (B) or 9 (C) days following the removal of all but three newly born nymphs. (B) Non-nicotine-tolerant *M. persicae* strain. (C) Nicotine-tolerant *M. persicae* strain. P values of student’s *t*-tests are displayed above mutants that were compared to wildtype, with a Benjamini-Hochberg correction. Solid lines represent median values. Dashed lines represent quartiles. (B) N = 8. (C) N = 9.

Whereas phloem feeders belong to the Hemiptera order, leaf chewers are found across many orders, including Lepidoptera, Coleoptera and Orthoptera. For this reason, in addition to two lepidopteran species, we examined a leaf chewing orthopteran, *Schistocerca americana* (American bird grasshopper). Four grasshoppers were confined to single wildtype, *a622*, *asat2* and *a622*+*asat2* plants and, after 16 days of feeding, were weighed and their survival rates were scored. Only grasshoppers confined to the double *a622*+*asat2* mutants were heavier than those confined to wildtype (Fig. 5A), and grasshoppers confined to all three mutants survived at higher rates compared to wildtype (Fig. 5B). When the effects of nicotine and acylsugars on grasshopper weight were examined in a two-way ANOVA, the effects of both nicotine (p=0.001) and acylsugars (p=0.002) were significant, whereas the interaction between them was not. In addition, choice assays between pairs of leaf discs of the four genotypes were performed (Fig. 5C-H). Grasshoppers fed more on *a622* and *a622*+*asat2* leaves compared to wildtype (Fig. 5C,E) and on *a622*+*asat2* leaves compared to *asat2* (Fig. 5H). The combination of these results indicates that both acylsugars and nicotine protect *N. benthamiana* from *S. americana* herbivory, but that nicotine has the greater effect of the two.

**Figure 5.**
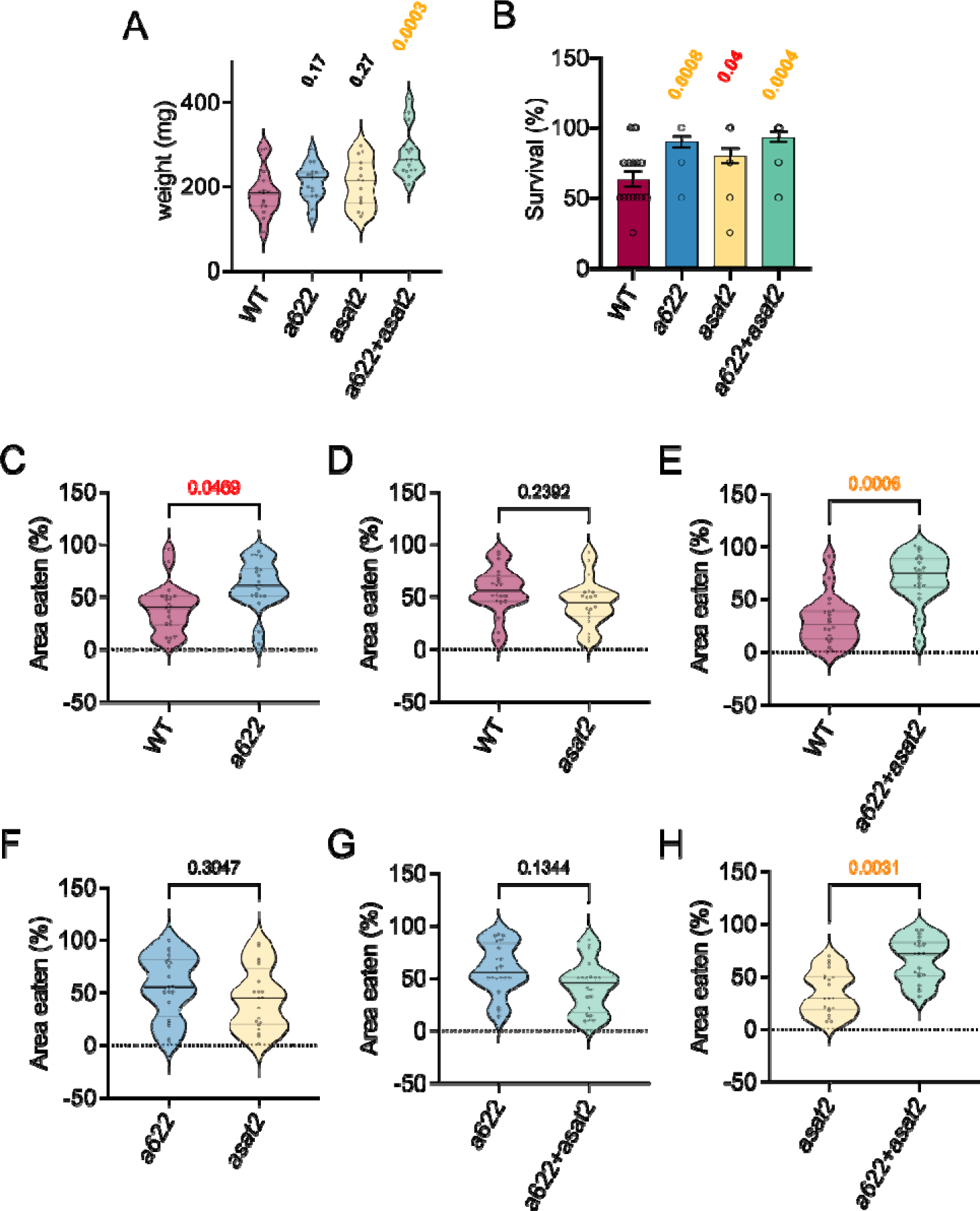
Growth, survival, and leaf choice of *Schistocerca americana* under controlled conditions. (A) Average weight and (B) percent survival of groups of four *S. americana* grasshoppers confined on whole wildtype and mutant *N. benthamiana* plants. N = 15; P values are from Benjamini Hochberg corrected student’s *t-*tests, comparing to the wildtype (WT) control. (C-H) Percentage of total consumed leaf area in plates with leaf discs from two different genotypes. The compared *N. benthamiana* lines are indicated beneath the panels. P values indicated above panels were obtained using paired Student’s *t*-tests. Solid and dashed lines represent median and quartile values, respectively. N = 20.

We next examined the effects of our mutant population on aphid reproduction using both nicotine-sensitive and nicotine-adapted *M. persicae* strains. Results were identical for the two strains, with aphids reproducing faster on mutants containing an *asat2* knockout (Fig. 4B-C). All other mutant combinations were similar to wildtype. When analyzing the effects of the compounds using a two-way ANOVA, similar to the results in *C. virescence*, only acyl sugars had a significant effect on aphid reproduction of both the non-adapted strain of *M, persicae* (p = 0.017) and the nicotine adapted strain (p = 0.0007). Just as when the plants were grown in the field, acylsugars dominated defense against aphids, and the other compounds were less important in this context. The lower protective capacity of nicotine was not related to the plants’ size in this case.

An additional question we wished to address, was whether aphids’ preference of plants harboring *asat2* mutations in the field was due to an increased “choice” of these plants, increased growth, reproduction and survival of the aphids, or a combination of the factors. We therefore performed dual choice assays using leaf discs. Twenty-one chemotype combinations were examined using the two aphid strains. We found that the nicotine-adapted strain, which did not show a different response compared to the nicotine sensitive strain in the reproduction assay, did not significantly prefer any of the chemotypes (Fig. S10). However, when all leaves that have an *asat2* mutation were compared against those that do not, a significant preference for the *asat2* mutants was revealed (Fig. S10W). Surprisingly, aphids from the “nicotine susceptible” strain did not show a preference for nicotine-free leaves (Fig. 6E, I, H, M, N, O, Q). Furthermore, when comparing all leaves containing normal nicotine abundance to those where nicotine was reduced, the aphids’ preference was similar (Fig. 6V). By contrast, all leaves harboring the *asat2* mutation were preferred compared to wildtype (Fig. 6B, D, F), and also relative to many of the other mutants (Fig. 6G, K, L, N, P, R, S, U). The combination of all *asat2* mutants compared to all leaf discs not harboring an *asat2* mutation, was significant as well (Fig. 6W).

**Figure 6.**
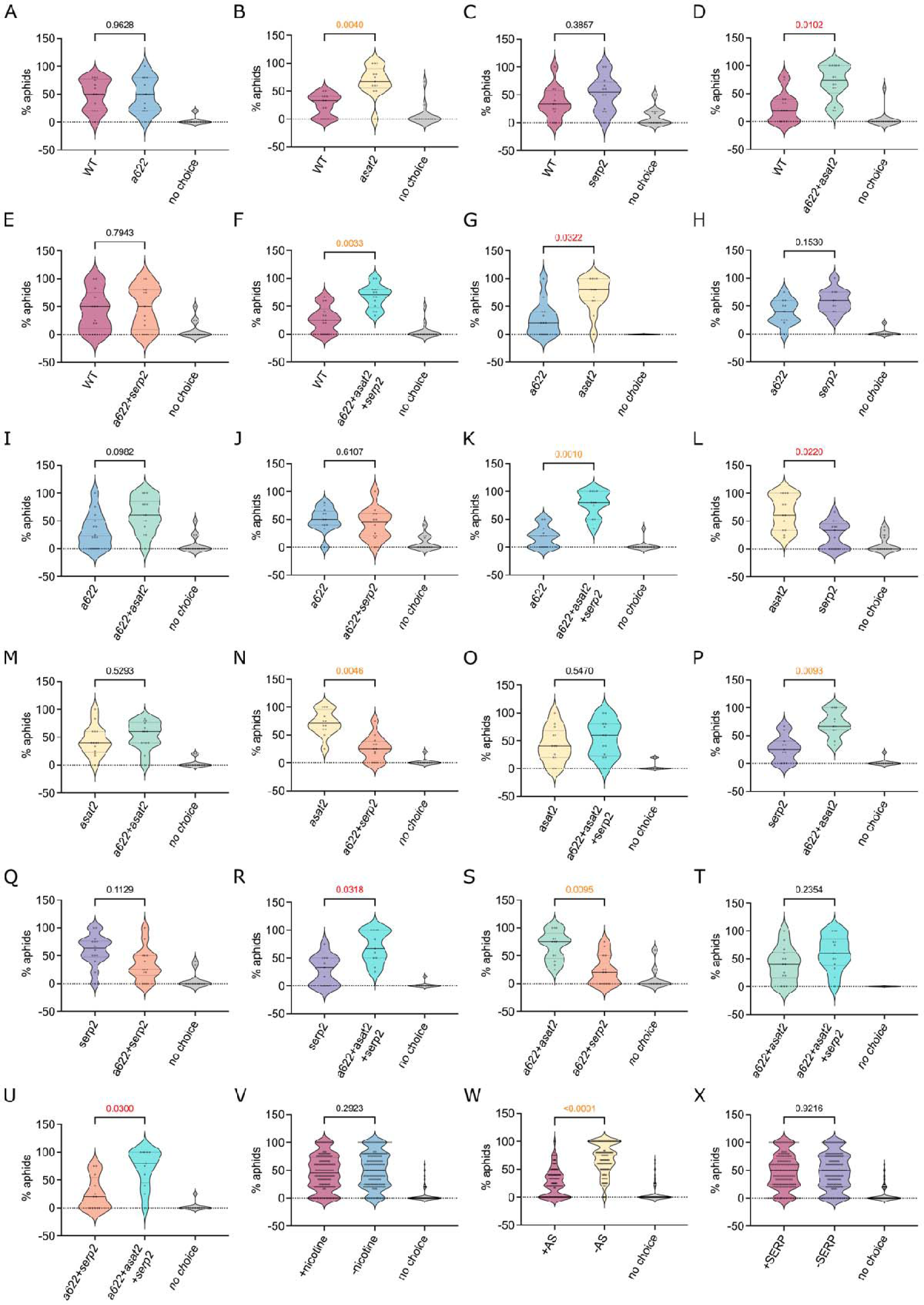
Non-nicotine-adapted *Myzus persicae* choice between 21 combinations of two leaf discs of wildtype and mutant plants. (A-U) comparison of single lines. The compared lines are indicated below the panels. (V-X) A comparison of all mutant discs harboring a mutation to all those that do not harbor it. P values indicated above panels were obtained from comparisons of the two-choice percentage using paired Student’s *t*-tests. Solid and dashed lines represent median and quartile values, respectively. (A-U) N = 11-14. (V-X) N = 137–152.

## Discussion

Chemical defenses are an extremely large group of plant protectants. A majority of these metabolites are assumed to be related to protection against pathogens and herbivores (Kortbeek *et al*., 2019). However, the effects of very few of these metabolites on insect interactions have been examined in detail using mutant lines. Furthermore, since each plant produces many specialized metabolites and other defenses simultaneously, these compounds may have an altered effect in the context of the entire plant’s chemotype.

For this reason, we chose three prominent *N. benthamiana* defensive compounds and examined their effects, both separately and combined. We found that, as might be expected, different defenses affected different interactors to varying extents. However, acylsugars stood out in almost all of the interaction types. Whether it was flea beetles, aphids, flies, or spiders, these sticky compounds protected plants from the initial visitation stage to the final colonization stage. Since the comparisons we performed were on plants that had a reduced abundance of the compounds, instances where only the double *a622*+*asat2* mutants were susceptible indicate that each compound on its own is enough to protect the plant. In this case, we might ask what advantage *N. benthamiana* gains from producing extremely high amounts of nicotine, which may reach up to 0.4% of the leaf’s dry weight (Drapal *et al*., 2021), which is similar to ∼0.02%-0.04% of leaf fresh weight reported for *N. attenuata* (Keinänen *et al*., 2001; Steppuhn *et al*., 2004). One probable answer, is that nicotine protects *N. benthamiana* from herbivores which were not represented in our field site. For instance, Machado et al., (Machado *et al*., 2016) found that jasmonate-deficient *Nicotiana attenuata* plants were more susceptible to vertebrate herbivores such as deer and rabbits. Our experimental setup, which excluded deer herbivory and attempted to exclude groundhog herbivory was not able to assess such an effect. Similarly, we found that nicotine has a greater effect than acylsugars on grasshopper herbivory under controlled conditions. However, no live grasshoppers were found in our field site. It is probable that, if our mutant population were grown in the native range of *S. americana*, greater effects of nicotine would be observed in the field.

It is possible that nicotine, like many other specialized metabolites (Zhou *et al*., 2018; Erb & Kliebenstein, 2020) has a developmental role, in addition to its insecticidal properties (Parr & Thurston, 1972; Steppuhn *et al*., 2004; Harvey *et al*., 2007). This is in line with the smaller size of our *N. benthamiana a622* mutants at lower temperatures, and a similar recent observation in *N. tabacum* mutants (Burner *et al*., 2022). By contrast targeting other parts of the nicotine biosynthesis pathway, RNAi of the putrescine methyl transferase (*PMT*) gene required for production of the methylpyrrolidine ring (Steppuhn *et al*., 2004) and genome editing of the berberine bridge like protein (*BBL*) gene (Vollheyde *et al*., 2023), did not inhibit growth of the nicotine-reduced plants. However, since the nicotine abundance in the *a622* mutants is reduced to a greater extent than in the *pmt* and *bbl* mutants, in addition to expected reduction in three other pyridine alkaloids, anabasine, anatabine, and nornicotine (Kaminski *et al*., 2020), it cannot be ruled out that pyridine alkaloids directly affect plant growth.

Nonetheless, the double *a622*+*asat2* mutants demonstrate that the small size of *a622* mutants is not the main reason for the fewer insect interactions observed on these plants. In all five interactions in which *asat2* mutants had an increased prevalence of invertebrates, the smaller double mutants displayed a significant effect as well. Furthermore, there are two interactions where the effects of double mutants were significant but those of single *asat2* mutants were not. When observing the plants in the field, there were many cases in which smaller plants harboring mutations in the *asat2* gene had far more interactors than nearby larger plants which did not.

Nicotine in *N. benthamiana* may also serve as a “second line of defense” against insect herbivory. Almost every potent specialized metabolite that evolved in plants has driven the evolution and adaptation of specialist herbivores, which are more tolerant or altogether resistant to the metabolite. Furthermore, even when not looking at specialists, there are entire classes of herbivores that could just be less susceptible to certain chemical defenses, as was seen with the snails and grasshoppers in this study and was reported for slug resistance to alkaloids (Self *et al*., 1964; Cornell & Hawkins, 2003; Aguiar & Wink, 2005; Ali & Agrawal, 2012; Altesor *et al*., 2014; Petschenka & Agrawal, 2015).

We may wonder, in that case, whether examining the effects of *N. benthamiana’s* defensive compounds so far away from its natural habitat, not only in terms of distance, but also in terms of the growing conditions of this desert-adapted plant, are relevant. Indeed, examining the ecology of *N. benthamiana* in such an ectopic setting may not shed light on many aspects of *N*. *benthamiana’s* ecology and native interactions with herbivores. However, it can elucidate the function of the examined metabolites in a more precise way. In *N, benthamiana’s* natural habitat, herbivores have been exposed and adapted to its defenses for millions of years. In this habitat, when an insect consumes a nicotine-rich leaf, we do not know whether this is because nicotine does not affect the insect or whether it is a specialist herbivore attracted by the presence of nicotine. In our conditions, we can be almost certain that the aphids that were found on the nicotine-containing plants, were not there because of acquired tolerance through exposure, but rather through more inherent properties of the compound’s interaction with this insect. Similarly, *N. benthamiana* is unlikely to suffer from slug consumption in its natural habitat. However, that does not reduce the significance of our finding that nicotine has little effect on slugs and snails.

Plants lacking nicotine, acylsugars, and the serine protease inhibitor are nevertheless lethal to *S. exigua* neonates, demonstrating that they must have additional potent protections. When analyzing the specialized metabolites present in *Nicotiana* species, Drapal *et al*. showed that a laboratory strain of *N. benthamiana* produced several pyridine alkaloids, chlorogenic acids, sterols, terpenes, hydrocarbons, and other specialized metabolites (Drapal *et al*., 2022). This wide array of chemicals is consistent with the observation that combinations of compounds are typically involved in protecting plants against any given herbivore (Chen, 2008). In contrast to *S. exigua*, snails, which caused immense damage in the field, consumed wildtype plants as readily as the mutant plants. We can therefore assume that, even if we didn’t see an effect of nicotine or SerPIN-II in the field, there are contexts in which these play a primary role in plant protection. Finally, it is possible that the combination of compounds is greater than the sum of their individual effects. This was elegantly demonstrated by Steppuhn et al. (Steppuhn & Baldwin, 2007), who showed that herbivores could compensate for a protease inhibitor by increasing the amount of plant material they consumed. However, nicotine prevented this compensation response. Such an effect can be seen in the caterpillar growth assays. Nicotine alone, acylsugars alone, and even the combination of the two does not affect tobacco budworm growth. However, adding the *serp2* mutations, which alone was not found to affect insect interactions in any of our assays, results in plants that are more susceptible to *C. virescens* (Fig. 4A). Similarly, *S. americana* nymphs grew faster than on wildtype plants only in the absence of both acylsugars and nicotine (Fig. 5A).

In this study, we adopted an approach of hypothesis-based experimentation in a natural setting (Negin & Aharoni, 2023). Using this approach, we recorded every interaction we were able to score, including flies that were resting on leaves for mere seconds, parasitized aphids, and bumblebees pollinating the *N. benthamiana* flowers. This type of experimentation is common practice in many ecological circles but is not often used when more “molecular” trials that include the generation of mutant populations are performed. An approach using mutant plants has the advantage of allowing us to determine the defensive functions of the metabolite combinations, while still leaving room for more complex answers beyond rejecting null hypothesis.

## Author Contributions

GJ planned, guided, and oversaw the project, performed experiments, analyzed data and revised the manuscript. BN planned the project, conducted the experiments, analyzed the data, and wrote the manuscript. FW conducted experiments, aided with insect identification and revised the manuscript. HDF assisted in insect identification, insect assay setup, and manuscript revisions.

## Acknowledgments

We thank Hojun Song for providing grasshopper eggs. This research was made possible by US Department of Agriculture postdoctoral fellowship number 2022-67012-36739 to BN, and by US Department of Agriculture award 2021-67013-33565 and a Triad Foundation award to GJ.

## References

Agrawal AA, Petschenka G, Bingham RA, Weber MG, Rasmann S. 2012. Toxic cardenolides: Chemical ecology and coevolution of specialized plant–herbivore interactions. New Phytologist 194: 28–45.

Aguiar R, Wink M. 2005. How do slugs cope with toxic alkaloids? CHEMOECOLOGY 15: 167–177.

Ali JG, Agrawal AA. 2012. Specialist versus generalist insect herbivores and plant defense. Trends in Plant Science 17: 293–302.

Altesor P, García Á, Font E, Rodríguez-Haralambides A, Vilaró F, Oesterheld M, Soler R, González A. 2014. Glycoalkaloids of wild and cultivated *Solanum*: Effects on specialist and generalist Insect herbivores. Journal of Chemical Ecology 40: 599–608.

Bally J, Jung H, Mortimer C, Naim F, Philips JG, Hellens R, Bombarely A, Goodin MM, Waterhouse PM. 2018. The rise and rise of *Nicotiana benthamiana*: A plant for all reasons. Annual Review of Phytopathology 56: 405–426.

Bally J, Nakasugi K, Jia F, Jung H, Ho SYW, Wong M, Paul CM, Naim F, Wood CC, Crowhurst RN, et al. 2015. The extremophile *Nicotiana benthamiana* has traded viral defence for early vigour. Nature Plants 1: 1–6.

Burner N, Kernodle SP, Steede T, Lewis RS. 2022. Editing of A622 genes results in ultra-low nicotine whole tobacco plants at the expense of dramatically reduced growth and development. Molecular Breeding 42: 20.

Carrillo L, Martinez M, Álvarez-Alfageme F, Castañera P, Smagghe G, Diaz I, Ortego F. 2011. A barley cysteine-proteinase inhibitor reduces the performance of two aphid species in artificial diets and transgenic Arabidopsis plants. Transgenic Research 20: 305–319.

Chen M-S. 2008. Inducible direct plant defense against insect herbivores: A review. Insect Science 15: 101–114.

Cornell HV, Hawkins BA. 2003. Herbivore responses to plant secondary compounds: A test of phytochemical coevolution theory. The American Naturalist 161: 507–522.

Deboer KD, Lye JC, Aitken CD, Su AK-K, Hamill JD. 2009. The *A622* gene in *Nicotiana glauca* (tree tobacco): Evidence for a functional role in pyridine alkaloid synthesis. Plant Molecular Biology 69: 299–312.

Demkura PV, Abdala G, Baldwin IT, Ballare CL. 2010. Jasmonate-dependent and -independent pathways mediate specific effects of solar ultraviolet B radiation on leaf phenolics and antiherbivore defense. Plant Physiology 152: 1084–1095.

Drapal M, Enfissi EMA, Fraser PD. 2021. Metabolic changes in leaves of N. tabacum and N. benthamiana during plant development. Journal of Plant Physiology 265: 153486.

Drapal M, Enfissi EMA, Fraser PD. 2022. The chemotype core collection of genus *Nicotiana*. The Plant Journal 110: 1516–1528.

Ellison EE, Nagalakshmi U, Gamo ME, Huang P, Dinesh-Kumar S, Voytas DF. 2020. Multiplexed heritable gene editing using RNA viruses and mobile single guide RNAs. Nature Plants 6: 620– 624.

Engler C, Kandzia R, Marillonnet S. 2008. A one pot, one step, precision cloning method with high throughput capability. PLOS ONE 3: e3647.

Erb M, Kliebenstein DJ. 2020. Plant secondary metabolites as defenses, regulators, and primary metabolites: The blurred functional trichotomy. Plant Physiology 184: 39–52.

Fan P, Leong BJ, Last RL. 2019. Tip of the trichome: Evolution of acylsugar metabolic diversity in Solanaceae. Current Opinion in Plant Biology 49: 8–16.

Fan P, Miller AM, Schilmiller AL, Liu X, Ofner I, Jones AD, Zamir D, Last RL. 2016. In vitro reconstruction and analysis of evolutionary variation of the tomato acylsucrose metabolic network. Proceedings of the National Academy of Sciences 113: E239–E248

Feng H, Acosta-Gamboa L, Kruse LH, Tracy JD, Chung SH, Nava Fereira AR, Shakir S, Xu H, Sunter G, Gore MA, et al. 2022. Acylsugars protect *Nicotiana benthamiana* against insect herbivory and desiccation. Plant Molecular Biology 109: 505–522.

Feng H, Jander G. 2023. Serine proteinase inhibitors from Nicotiana benthamiana, a non-preferred host plant, inhibit the growth of Myzus persicae (green peach aphid). BioRxiv: 2023.05.16.540980.

Fobes JF, Mudd JB, Marsden MP. 1985. Epicuticular lipid accumulation on the leaves of *Lycopersicon pennellii* (Corr.) D’Arcy and *Lycopersicon esculentum* Mill. Plant Physiology 77: 567–570.

Green TR, Ryan CA. 1972. Wound-induced proteinase inhibitor in plant leaves: A possible defense mechanism against insects. Science 175: 776–777.

Hare JD. 2005. Biological activity of acyl glucose esters from *Datura wrightii* glandular trichomes against three native insect herbivores. Journal of Chemical Ecology 31: 1475–1491.

Hartl M, Giri AP, Kaur H, Baldwin IT. 2010. Serine protease inhibitors specifically defend *Solanum nigrum* against generalist herbivores but do not influence plant growth and development. The Plant Cell 22: 4158–4175.

Harvey JA, Van Dam NM, Witjes LMA, Soler R, Gols R. 2007. Effects of dietary nicotine on the development of an insect herbivore, its parasitoid and secondary hyperparasitoid over four trophic levels. Ecological Entomology 32: 15–23.

Heiling S, Li J, Halitschke R, Paetz C, Baldwin IT. 2022. The downside of metabolic diversity: Postingestive rearrangements by specialized insects. Proceedings of the National Academy of Sciences 119: e2122808119.

Hoogshagen M, Hastings AP, Chavez J, Duckett M, Pettit R, Pahnke AP, Agrawal AA, de Roode JC. 2023. Mixtures of milkweed cardenolides protect monarch butterflies against parasites. Journal of Chemical Ecology.

Hopkins RJ, van Dam NM, van Loon JJA. 2009. Role of glucosinolates in insect-plant relationships and multitrophic interactions. Annual Review of Entomology 54: 57–83.

Hu L, Ye M, Kuai P, Ye M, Erb M, Lou Y. 2018. OsLRR-RLK1, an early responsive leucine-rich repeat receptor-like kinase, initiates rice defense responses against a chewing herbivore. New Phytologist 219: 1097–1111.

Jassbi AR, Zare S, Asadollahi M, Schuman MC. 2017. Ecological roles and biological activities of specialized metabolites from the genus *Nicotiana*. Chemical Reviews 117: 12227–12280.

Kaminski KP, Goepfert S, Ivanov NV, Peitsch MC. 2020. Production of valuable compounds in tobacco. In: Ivanov NV, Sierro N, Peitsch MC, eds. Compendium of Plant Genomes. The Tobacco Plant Genome. Cham: Springer International Publishing, 249–263.

Keinänen M, Oldham NJ, Baldwin IT. 2001. Rapid HPLC screening of jasmonate-induced increases in tobacco alkaloids, phenolics, and diterpene glycosides in *Nicotiana attenuata*. Journal of Agricultural and Food Chemistry 49: 3553–3558.

Kortbeek RWJ, van der Gragt M, Bleeker PM. 2019. Endogenous plant metabolites against insects. European Journal of Plant Pathology 154: 67–90.

Kroumova AB, Wagner GJ. 2003. Different elongation pathways in the biosynthesis of acyl groups of trichome exudate sugar esters from various solanaceous plants. Planta 216: 1013– 1021.

Kumar P, Pandit SS, Steppuhn A, Baldwin IT. 2014. Natural history-driven, plant-mediated RNAi-based study reveals CYP6B46’s role in a nicotine-mediated antipredator herbivore defense. Proceedings of the National Academy of Sciences 111: 1245–1252.

Lei Y, Lu L, Liu H-Y, Li S, Xing F, Chen L-L. 2014. CRISPR-P: A web tool for synthetic single-guide RNA design of CRISPR-system in plants. Molecular plant 7: 1494–1496.

Li J-F, Norville JE, Aach J, McCormack M, Zhang D, Bush J, Church GM, Sheen J. 2013. Multiplex and homologous recombination–mediated genome editing in Arabidopsis and *Nicotiana benthamiana* using guide RNA and Cas9. Nature Biotechnology 31: 688–691.

Lou Y-R, Anthony TM, Fiesel PD, Arking RE, Christensen EM, Jones AD, Last RL. 2021. It happened again: Convergent evolution of acylglucose specialized metabolism in black nightshade and wild tomato. Science Advances 7: eabj8726.

Luu VT, Weinhold A, Ullah C, Dressel S, Schoettner M, Gase K, Gaquerel E, Xu S, Baldwin IT. 2017. O-acyl sugars protect a wild tobacco from both native fungal pathogens and a specialist herbivore. Plant Physiology 174: 370–386.

Machado RA, McClure M, Hervé MR, Baldwin IT, Erb M. 2016. Benefits of jasmonate-dependent defenses against vertebrate herbivores in nature. eLife 5: e13720.

Meihls LN, Handrick V, Glauser G, Barbier H, Kaur H, Haribal MM, Lipka AE, Gershenzon J, Buckler ES, Erb M, et al. 2013. Natural variation in maize aphid resistance is associated with 2,4-dihydroxy-7-methoxy-1,4-benzoxazin-3-one glucoside methyltransferase activity. The Plant Cell 25: 2341–2355.

Millar NS, Denholm I. 2007. Nicotinic acetylcholine receptors: Targets for commercially important insecticides. Invertebrate Neuroscience 7: 53–66.

Negin B, Aharoni A. 2023. Let’s connect nature with hypothesis-based experimentation and explore life in context. The Plant Journal 113: 23–25.

Negin B, Jander G. 2023. Convergent and divergent evolution of plant chemical defenses. Current Opinion in Plant Biology 73: 102368.

Negin B, Shachar L, Meir S, Ramirez CC, Rami Horowitz A, Jander G, Aharoni A. 2024. Fatty alcohols, a minor component of the tree tobacco surface wax, are associated with defence against caterpillar herbivory. Plant, Cell & Environment 47: 664–681.

Parr JC, Thurston R. 1972. Toxicity of nicotine in synthetic diets to larvae of the tobacco hornworm. Annals of the Entomological Society of America 65: 1185–1188.

Petschenka G, Agrawal AA. 2015. Milkweed butterfly resistance to plant toxins is linked to sequestration, not coping with a toxic diet. Proceedings of the Royal Society B: Biological Sciences 282: 20151865.

Pichersky E, Lewinsohn E. 2011. Convergent evolution in plant specialized metabolism. Annual Review of Plant Biology 62: 549–566.

Puterka GJ, Farone W, Palmer T, Barrington A. 2003. Structure-function relationships affecting the insecticidal and miticidal activity of sugar esters. Journal of Economic Entomology 96: 636– 644.

Ramsey JS, Elzinga DA, Sarkar P, Xin Y-R, Ghanim M, Jander G. 2014. Adaptation to nicotine feeding in *Myzus persicae*. Journal of Chemical Ecology 40: 869–877.

Ramsey JS, Wilson AC, de Vos M, Sun Q, Tamborindeguy C, Winfield A, Malloch G, Smith DM, Fenton B, Gray SM, et al. 2007. Genomic resources for Myzus persicae: EST sequencing, SNP identification, and microarray design. BMC Genomics 8: 423.

Reichstein T, von Euw J, Parsons JA, Rothschild M. 1968. Heart poisons in the monarch butterfly. Science 161: 861–866.

Schilmiller AL, Charbonneau AL, Last RL. 2012. Identification of a BAHD acetyltransferase that produces protective acyl sugars in tomato trichomes. Proceedings of the National Academy of Sciences 109: 16377–16382.

Schilmiller AL, Gilgallon K, Ghosh B, Jones AD, Last RL. 2016. Acylsugar acylhydrolases: carboxylesterase-catalyzed hydrolysis of acylsugars in tomato trichomes. Plant Physiology 170: 1331–1344.

Self LS, Guthrie FE, Hodgson E. 1964. Adaptation of tobacco hornworms to the ingestion of nicotine. Journal of Insect Physiology 10: 907–914.

Sparkes IA, Runions J, Kearns A, Hawes C. 2006. Rapid, transient expression of fluorescent fusion proteins in tobacco plants and generation of stably transformed plants. Nature Protocols 1: 2019–2025.

Steppuhn A, Baldwin IT. 2007. Resistance management in a native plant: Nicotine prevents herbivores from compensating for plant protease inhibitors. Ecology Letters 10: 499–511.

Steppuhn A, Gase K, Krock B, Halitschke R, Baldwin IT. 2004. Nicotine’s defensive function in nature. PLOS Biology 2: e217.

Van Dam NM, Hare JD. 1998. Biological activity of *Datura wrightii* glandular trichome exudate against *Manduca sexta* larvae. Journal of Chemical Ecology 24: 1529–1549.

Van Dam NM, Horn M, Mareš M, Baldwin IT. 2001. Ontogeny constrains systemic protease inhibitor response in *Nicotiana attenuata*. Journal of Chemical Ecology 27: 547–568.

Vollheyde K, Dudley QM, Yang T, Oz MT, Mancinotti D, Fedi MO, Heavens D, Linsmith G, Chhetry M, Smedley MA, et al. 2023. An improved *Nicotiana benthamiana* bioproduction chassis provides novel insights into nicotine biosynthesis. New Phytologist 240: 302–317.

Wang S, Alseekh S, Fernie AR, Luo J. 2019. The structure and function of major plant metabolite modifications. Molecular Plant 12: 899–919.

Xu S, Brockmöller T, Navarro-Quezada A, Kuhl H, Gase K, Ling Z, Zhou W, Kreitzer C, Stanke M, Tang H, et al. 2017. Wild tobacco genomes reveal the evolution of nicotine biosynthesis. Proceedings of the National Academy of Sciences 114: 6133–6138.

Yang RSH, Guthrie FE. 1969. Physiological responses of insects to nicotine. Annals of the Entomological Society of America 62: 141–146.

Yang C, Halitschke R, O’Connor SE. 2023. OXIDOSQUALENE CYCLASE 1 and 2 influence triterpene biosynthesis and defense in *Nicotiana attenuata*. Plant Physiology.

Zavala JA, Patankar AG, Gase K, Hui D, Baldwin IT. 2004. Manipulation of endogenous trypsin proteinase inhibitor production in *Nicotiana attenuata* demonstrates their function as antiherbivore defenses. Plant Physiology 134: 1181–1190.

Zhou S, Richter A, Jander G. 2018. Beyond defense: Multiple functions of benzoxazinoids in maize metabolism. Plant and Cell Physiology 59: 1528–1537.

Züst T, Strickler SR, Powell AF, Mabry ME, An H, Mirzaei M, York T, Holland CK, Kumar P, Erb M. 2020. Independent evolution of ancestral and novel defenses in a genus of toxic plants (Erysimum, Brassicaceae). Elife 9.

