## Supplementary figures S1-S10 and table S1 for "Knockout mutations of *Nicotiana benthamiana* defenses reveal the relative importance of acylsugars, nicotine, and a serine protease inhibitor in a natural setting"

**This file includes:**

Figures S1 to S10

Table S1

Legend for Datasets S1-S3

**Other supporting materials for this manuscript include the following:**

Dataset S1

Dataset S2

Dataset S3

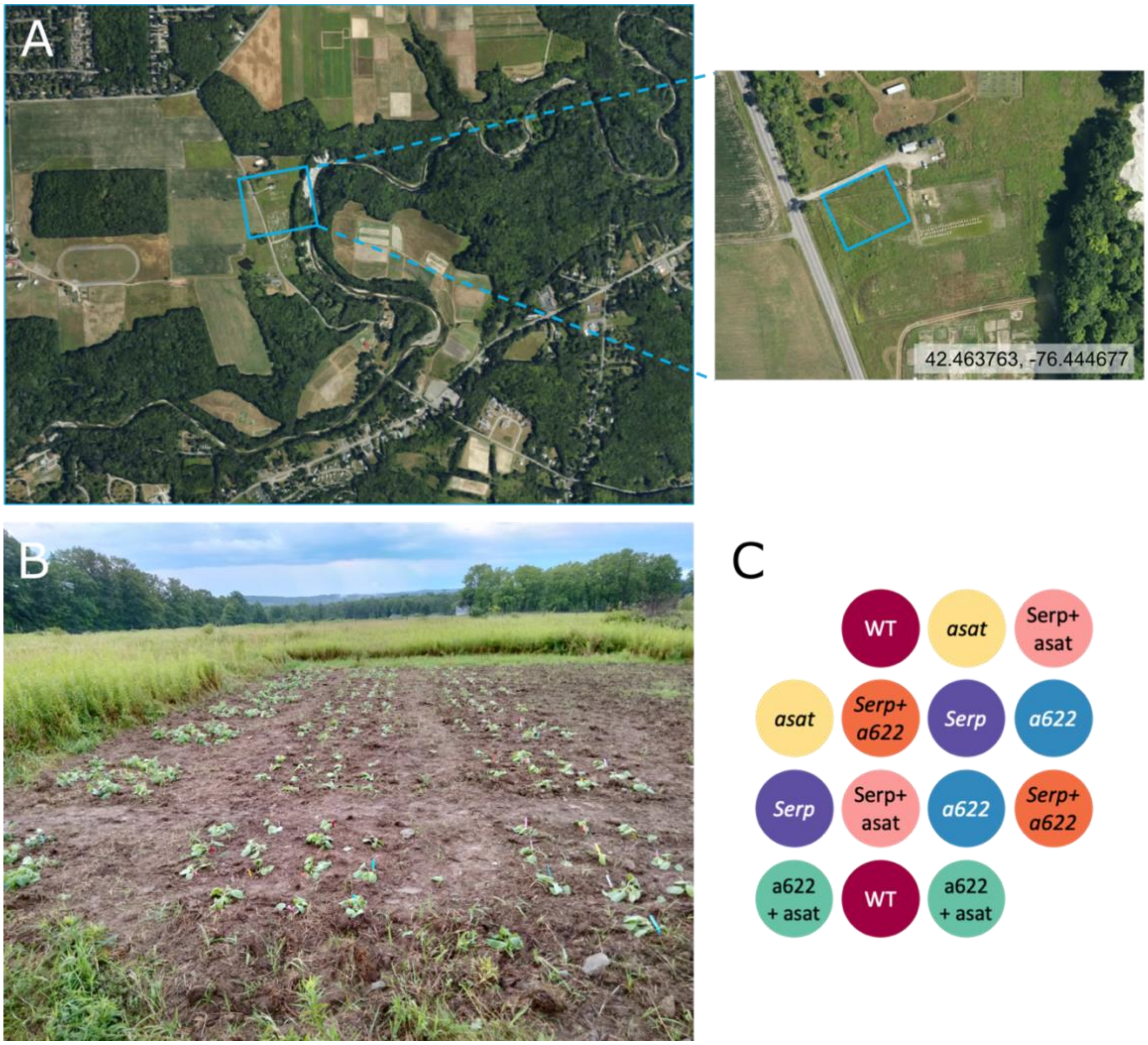


**Fig. S1.** Field location and setup. (A**)** A satellite image of the field demonstrates its location in the midst of both agricultural and natural areas. The field itself was a natural area that was deer-excluded by a fence and plowed in preparation for planting. (B) The field immediately following the planting of the 24 blocks of 14 plants. Golden rod (*Solidago* genus) can be seen dominating the unplowed field margins. (C) A representative plant setup in one of the blocks.

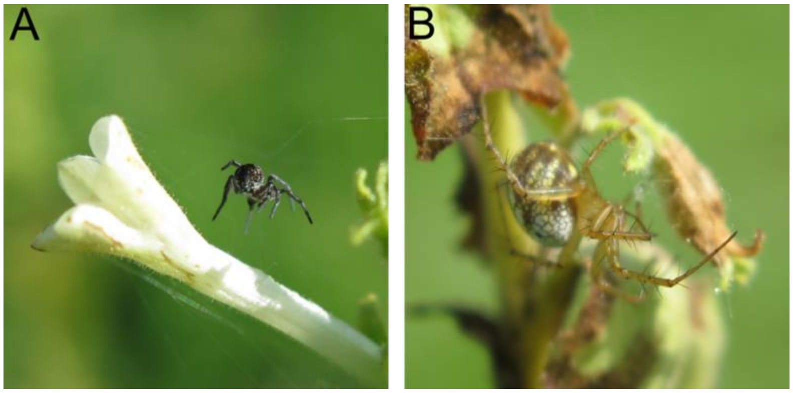


**Fig. S2**. Photographs of spiders found in consecutive observations on the same plant, an *asat2* mutant positioned at block 18, plant 7. (A) A spider possibly from the *Araneus* genus. (B) An orb weaver spider, probably *Mangora acalypha*.

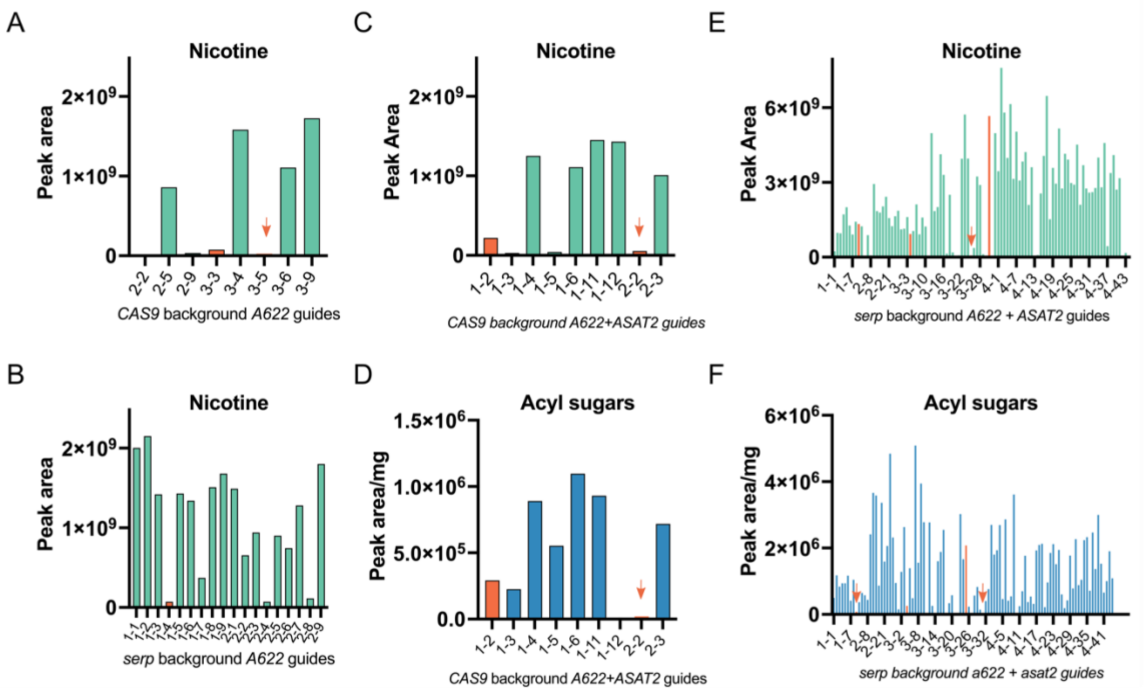


**Fig. S3.** Screening of nicotine and acylsugar abundance in T1 *a622* and *asat2* mutants. Both metabolites were extracted from leaves and relatively quantified using liquid chromatography-mass spectrometry. Lines that were selected to continue to T2 analysis or whose progeny were used directly are indicated by orange bars, or arrows when bars are not visible. (A**)** Nicotine abundance of *a622* mutants; the *3-3* and *3-5* lines were selected. (B) Nicotine abundance of *a622* mutants in the *serp2* background; the *1-4* line was selected. (C) Nicotine abundance of *a622+asat2* double mutants; the *1-2* and *2-2* lines were selected. (D) Acylsugar abundance of *a622+asat2* double mutants, showing the selected lines. (E) Nicotine abundance and (F) acylsugar abundance of *a622+asat2+serp2* mutant candidates. The *1-9* and *3-31* lines were selected as *a622+serp2* mutants. The *3-25* line was selected as an *a622+serp2* mutant. The *3-4* line, which has a mutation in the *ASAT2* gene and a heterozygous mutation in the *A622* gene, was selected to screen for triple mutants in the next generation. Bars represent individual plants' values.

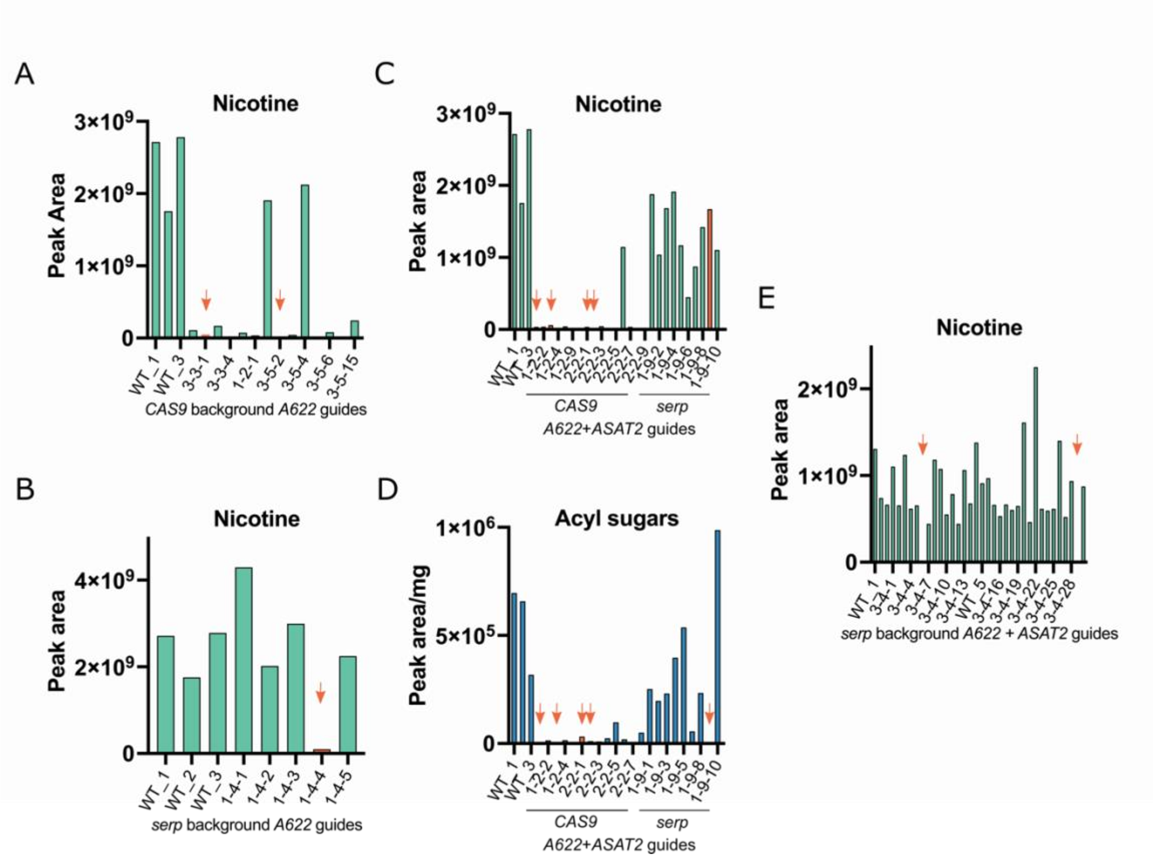


**Fig. S4.** Screening of nicotine and acylsugar abundance in T2 *a622* and *asat2* mutants. Both metabolites were extracted from leaves and relatively quantified using liquid chromatography-mass spectrometry. Lines that were selected are indicated by orange bars, or arrows when bars are not visible. (A) Nicotine abundance of T2 *a622* mutants; the *3-3-1* and *3-5-2* lines were selected. (B) Nicotine abundance of *a622* mutants in the *serp2* background; the *1-4-4* line was selected. (C) Nicotine abundance of *a622+asat2* double mutants, and *asat2+serp2* mutants; the *1-2-1, 1-2-3, 2-2-1* and *2-2-2* lines were selected as *a622+asat2* mutants, and the *1-9-9* line was selected as an *asat2+serp2* mutant. (D) Acylsugar relative abundance of the same mutant set as in panel C. The same mutant lines are indicated. (E) Nicotine abundance of the *serp2* background *3-4* lines. The *3-4-6* and *3-4-29* lines were selected as triple mutants. Bars represent individual plants' values.

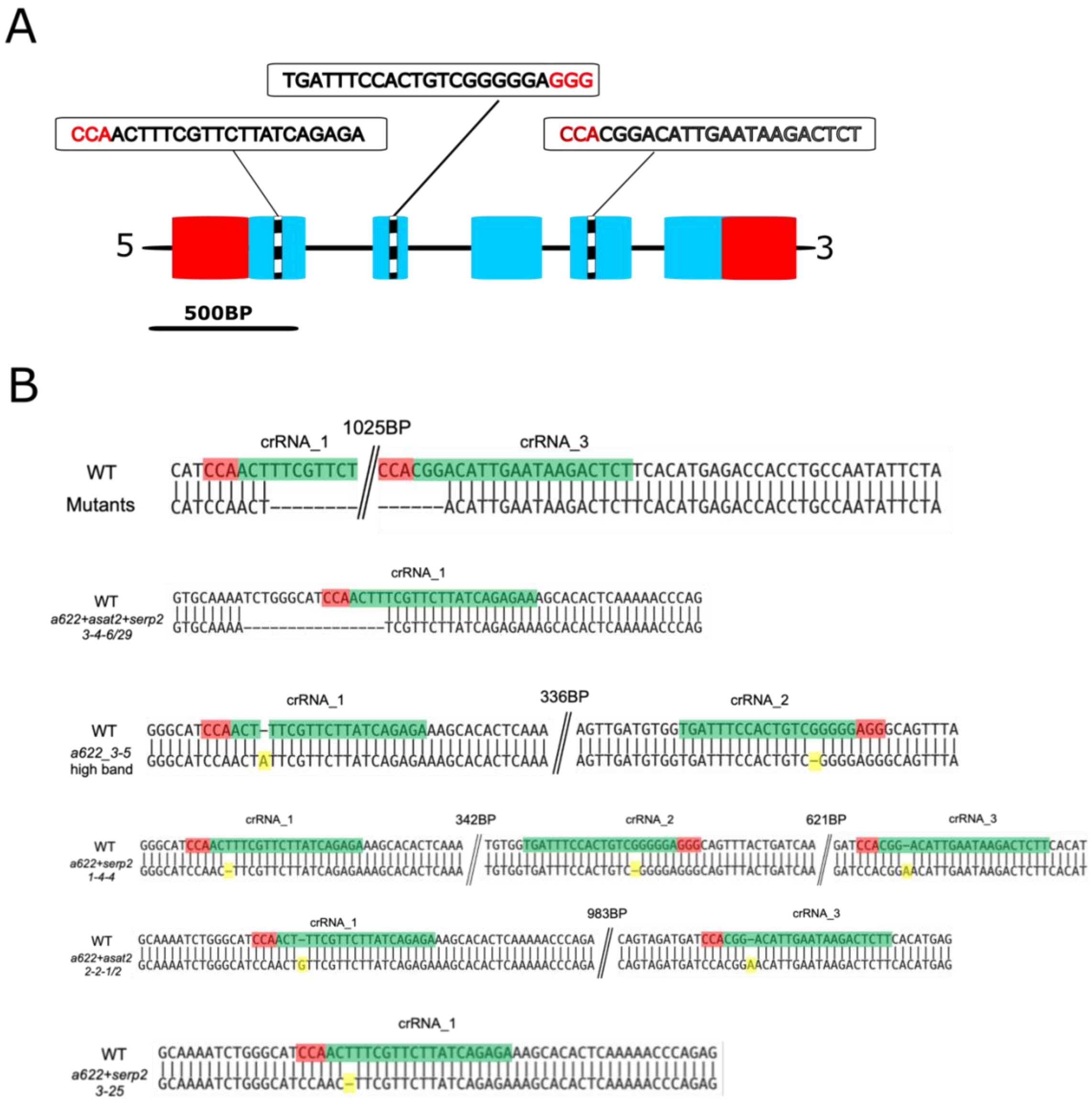


**Fig. S5.** Characterization of CRISPR-CAS9-derived genome edits in the *A622* gene in the mutant population used in this study. (A) A scheme of the *A622* gene targeted for genome editing in this study. Red boxes represent UTRs, blue boxes represent exons, and the black line between boxes represents introns. The region of the crRNA is indicated by black and white boxes (not proportional to the crRNA's actual 20 BP size). (B) Different mutations in the *A622* gene of the mutant population. Red coloring indicates the PAM sequence, green coloring the crRNA sequence and yellow coloring highlights single base indels. The topmost 1039BP deletion between crRNA 1-3 was present in several lines (see Table S1).

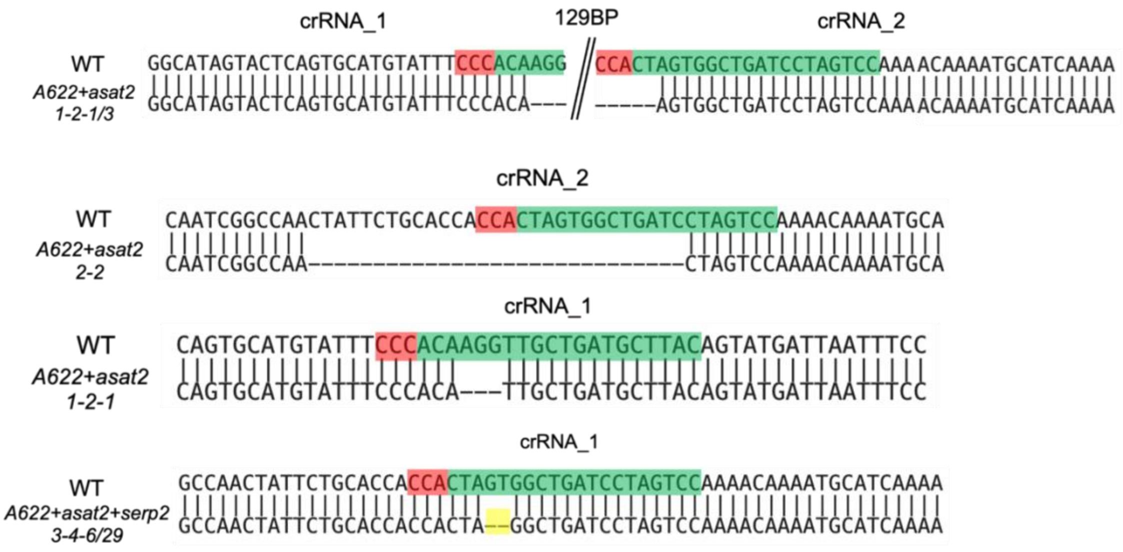


**Fig. S6.** Characterization of CRISPR-CAS9 derived genome edits in the *ASAT2* gene in the mutant population used in this study. Different genome edits created in this study are presented. Red coloring indicates the PAM sequence, green coloring the crRNA sequence, and yellow coloring highlights a 2BP deletion.

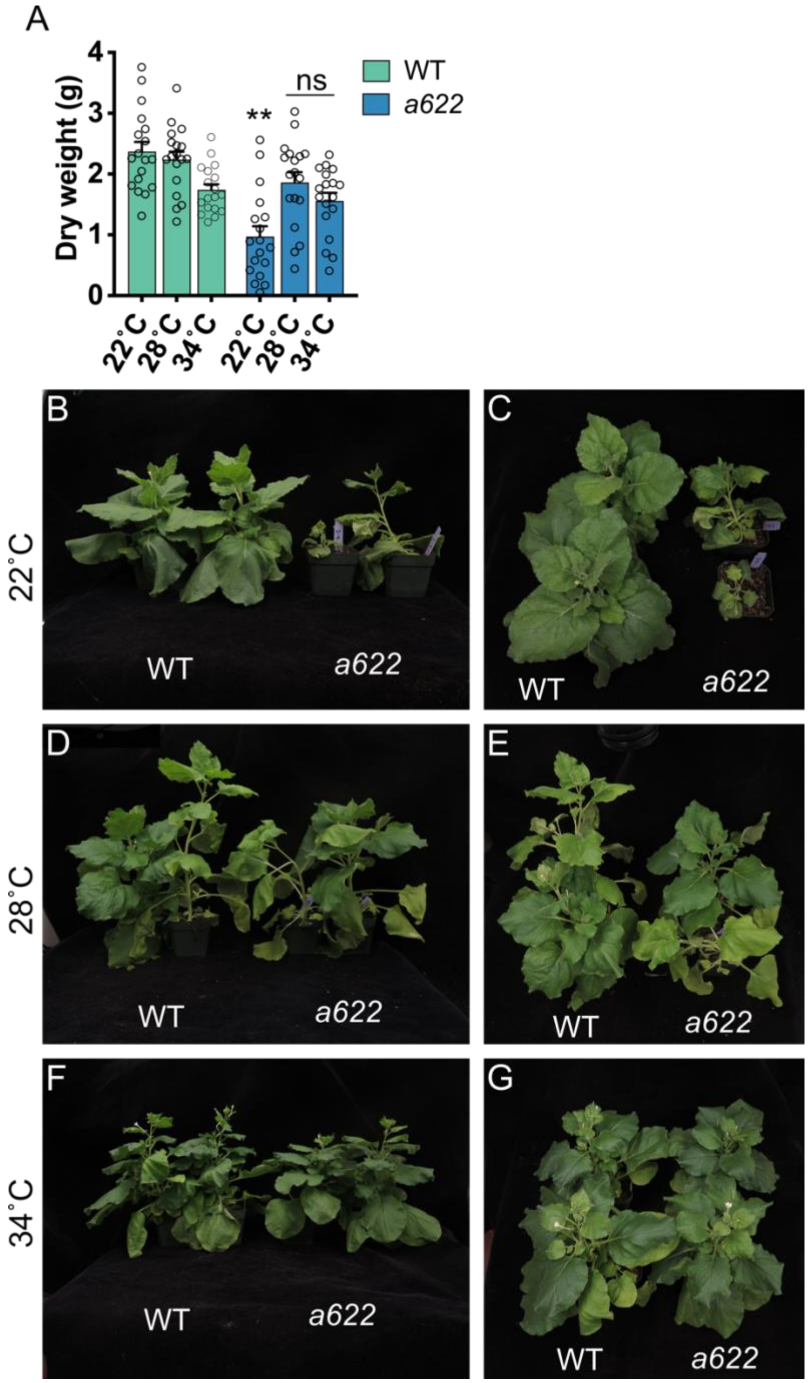


**Fig. S7.** Effects of growth temp. on plant size of wildtype and *a622* mutants. (A) Dry weight of four-week-old wildtype and *a622* mutants grown in a growth room at 22˚C, or in growth chambers at 28˚C or 34˚C. Double asterisks represent significance of $p < 0.01$. N = 18. (B-G) Photographs of representative plants grown in the different conditions. (B-C) 22˚C. (D-E) 28˚C. (F-G) 34˚C. In each photograph, two wildtype plants are displayed on the left-hand side beside two *a622* mutants.

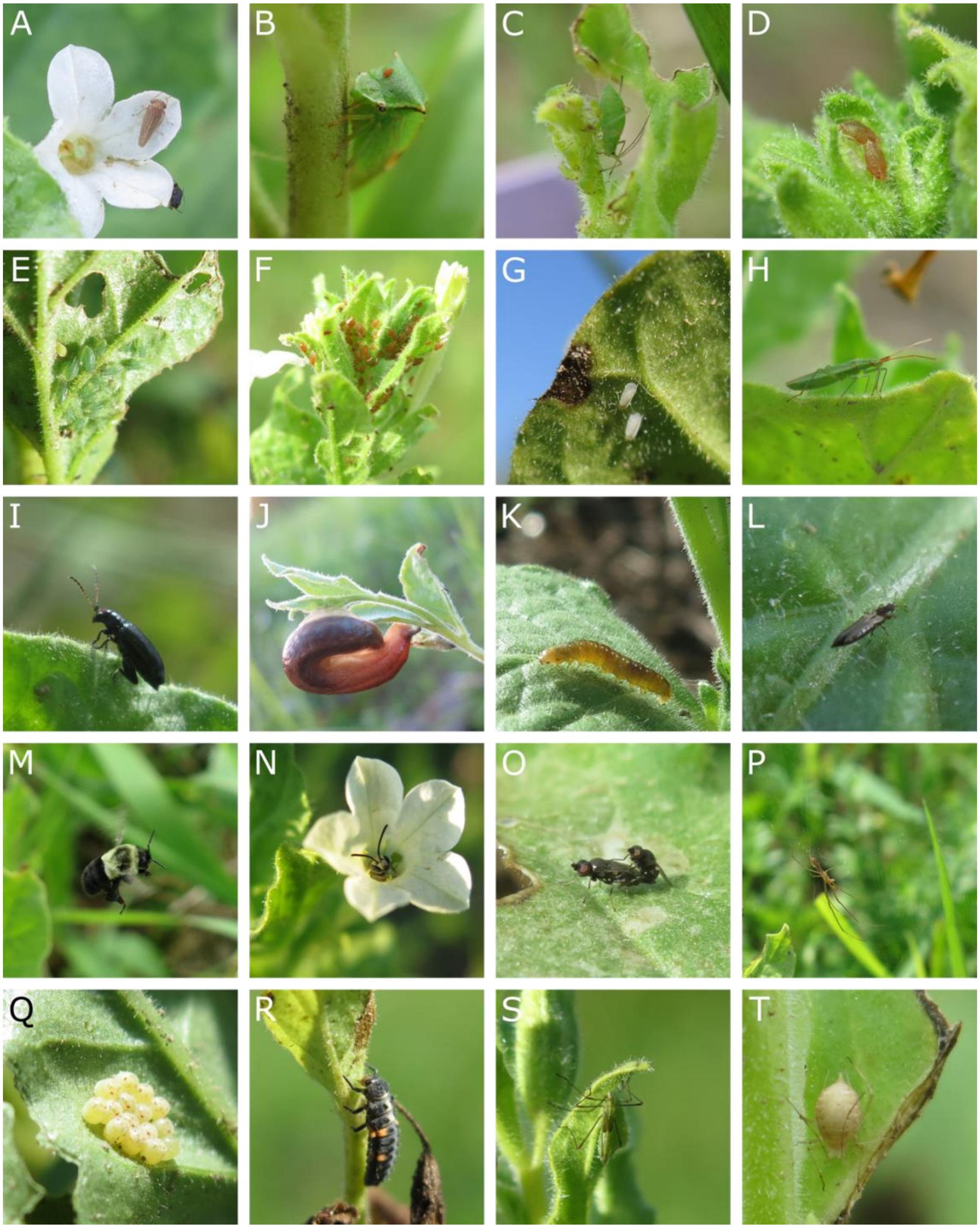

**Fig. S8.** Additional invertebrates observed on field grown *Nicotiana benthamiana*. (A-H) Insects from the phloem feeding guild. (I-K) Leaf chewers. (A) A leafhopper from the *Agallia* genus. (B) A buffalo tree hopper (*Stictocephala bisonia*). (C) A leaf aphid. (D) A leaf aphid, probably *Myzus persicae.* (E-F) Leaf aphid colonies. (G) Whiteflies. (H) A rice leaf bug (*Trigonotylus caelestialium*). (I**)** A flea beetle from the *Epitrix* genus. (J**)** A slug, possibly from the species *Arion subfuscus*. (K) Possibly an *Ancylis muricana* caterpillar, the only caterpillar found on plants throughout the experiment. (L) A thrips, probably *Heliothrips haemorrhoidalis*. (M) A bumble bee (genus *Bombus*) at flight while pollinating *N. benthamiana.* (N) A solitary bee resting in an *N. benthamiana* flower. (O) Flies mating on the leaf of an *a622+serp2* mutant. (P) A Long-jawed orb weaver spider (*Tetragnatha montana*). (Q) *Halyomorpha halys* eggs. (R) A ladybug (*Coccinella septempunctata*) larva. (S) An assassin bug, possibly from the genus *Zelus*. (T) An aphid parasitized by a wasp larva.

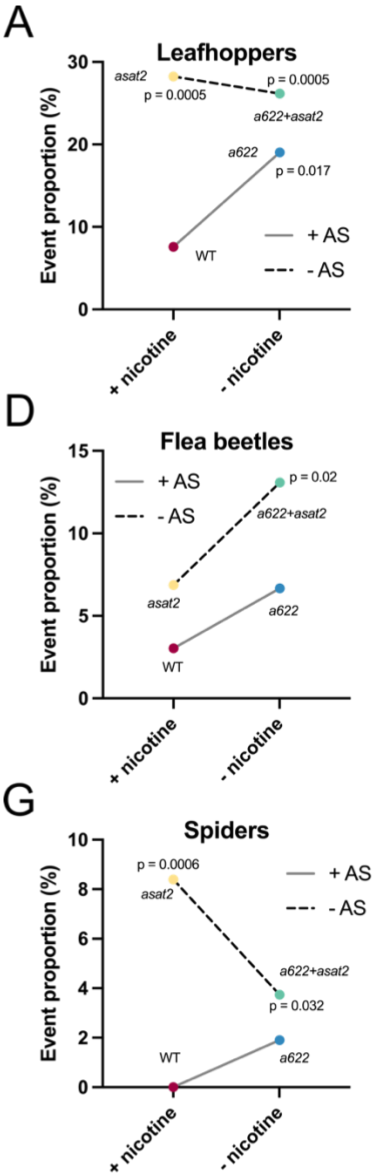


**Fig. S9.** Interactions between nicotine and acylsugar mutations and invertebrate interactions. The additive effect of loss of both nicotine and acylsugars may be seen in panel (D), whereas in other panels the loss of nicotine does not make the plants more susceptible to invertebrates. Only panels where a significant or marginally significant effect of acylsugars or nicotine were found compared to wildtype are displayed.

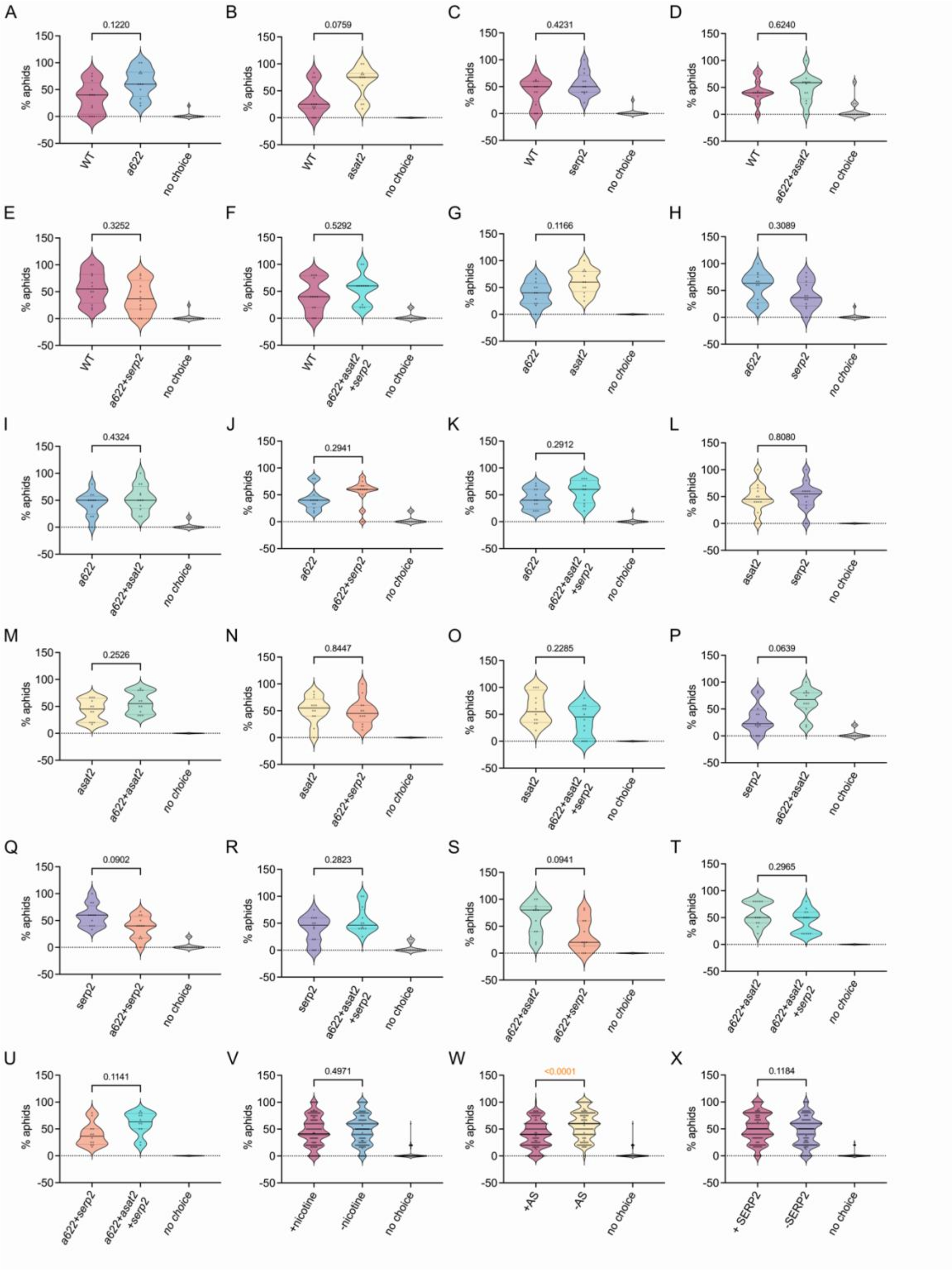
A
0.1220
% aphids
WT
a622
no choice
B
0.0759
% aphids
WT
asat2
no choice
C
0.4231
% aphids
WT
serp2
no choice
D
0.6240
% aphids
WT
a622+asat2
no choice
E
0.3252
% aphids
WT
a622+serp2
no choice
F
0.5292
% aphids
WT
a622+asat2 +serp2
no choice
G
0.1166
% aphids
a622
asat2
no choice
H
0.3089
% aphids
a622
serp2
no choice
I
0.4324
% aphids
a622
a622+asat2
no choice
J
0.2941
% aphids
a622
a622+serp2
no choice
K
0.2912
% aphids
a622
a622+asat2 +serp2
no choice
L
0.8080
% aphids
asat2
serp2
no choice
M
0.2526
% aphids
asat2
a622+asat2
no choice
N
0.8447
% aphids
asat2
a622+serp2
no choice
O
0.2285
% aphids
asat2
a622+asat2 +serp2
no choice
P
0.0639
% aphids
serp2
a622+asat2
no choice
Q
0.0902
% aphids
serp2
a622+serp2
no choice
R
0.2823
% aphids
serp2
a622+asat2 +serp2
no choice
S
0.0941
% aphids
a622+asat2
a622+serp2
no choice
T
0.2965
% aphids
a622+asat2
a622+asat2 +serp2
no choice
U
0.1141
% aphids
a622+serp2
a622+asat2 +serp2
no choice
V
0.4971
% aphids
+nicotine
-nicotine
no choice
W
<0.0001
% aphids
+AS
-AS
no choice
X
0.1184
% aphids
+ SERP2
-SERP2
no choice

**Fig. S10.** Nicotine-tolerant *Myzus persicae* choice between 21 combinations of two leaf discs of wildtype and mutant plants. (A-U) comparison of single lines. The compared lines are indicated below the panels. (V-X) A comparison of all mutant discs harboring a specific mutation to all those that do not harbor it. P values are indicated above panels and were obtained from comparisons of the two-choice percentage using a paired Student's *t*-test. Solid lines represent median values. Dashed lines represent quartiles. (A-U) N = 12. (V-X) N = 144.

**Table S1.** A summary of the different CRISPR induced mutations in the *N. benthamiana* mutant population

| Mutations | line | *A622* | *ASAT2* | *SERP2* |
|---|---|---|---|---|
| *asat2* | *asat2-1* | - | "A" deletion in the first target sequence | - |
| *asat2* | *asat2-2* | - | 118 BP deletion, with a 4BP insertion in it | - |
| *serp2* | *Serp2-1* | - | - | "T" insertion in target sequence 2 |
| *serp2* | *Serp2-2* | - | - | 3BP deletion in target sequence 2 |
| *a622* | 3-3-1 | 1039BP deletion between target sequences 1-3 | - | - |
| *a622* | 3-5-2 | "A" insertion in target sequence 1, "G" deletion in target sequence 2 // Deletion between target sequences 1-3 | - | - |
| *a622 + asat2* | 1-2-1 | 1039BP deletion between target sequences 1-3 | 136BP deletion between target sequences 1-2, 3 BP deletion in target sequence 1 | - |
| *a622 + asat2* | 1-2-3 | 1039BP deletion between target sequences 1-3 | 136BP deletion between target sequences 1-2 | - |
| *a622 + asat2* | 2-2-1 | "G" insertion in target sequence 1, "A" insertion in target sequence 3 | 136BP deletion between target sequences 1-2, 29 BP deletion in target sequence 2 | - |
| *a622 + asat2* | 2-2-2 | "G" insertion in target sequence 1 | 136BP deletion between target sequences 1-2, 29 BP deletion in target sequence 2 | - |
| *a622 + serp2* | 1-4-4 | "T" deletion in target sequence 1, "G" deletion in target sequence 2, "A" insertion in target sequence 3 | - | 3BP deletion in target sequence 2 |
| *a622 + serp2* | 3-25 | "T" deletion in target sequence 1 | - | 3BP deletion in target sequence 2 |
| *Serp2* | 1-9-9 | - | - | 3BP deletion in target sequence 2 |
| *Serp2* | 3-31 | - | - | 3BP deletion in target sequence 2 |
| *a622 + asat2 + serp2* | 3-4-6 | 16BP deletion in target sequence 1 | "GT" deletion in target sequence 2 | 3BP deletion in target sequence 2 |
| *a622 + asat2 + serp2* | 3-4-29 | 16BP deletion in target sequence 1 | "GT" deletion in target sequence 2 | 3BP deletion in target sequence 2 |

.

**Dataset S1 (separate file).** Observations of *Nicotiana benthamiana* field grown plants and invertebrate interactions.

**Dataset S2 (separate file).** Observations of invertebrate interactions of select invertebrates (leaf hoppers, aphid colonies, flea beetles and spiders), divided by plant throughout the observation period.

**Dataset S3 (separate file).** The genomic sequence of the *A622* gene targeted for genome editing in this study. Green coloring represents exons. crRNAs are indicated in green. PAMs, ATG and stop codons are indicated in red.
